# A Purkinje cell to parabrachial nucleus pathway enables broad cerebellar influence over the forebrain

**DOI:** 10.1101/2021.09.21.461236

**Authors:** Christopher H. Chen, Leannah N. Newman, Amanda P. Stark, Katherine E. Bond, Dawei Zhang, Stefano Nardone, Charles R. Vanderburg, Naeem M. Nadaf, Zhiyi Yao, Kefiloe Mutume, Isabella Flaquer, Bradford B. Lowell, Evan Z. Macosko, Wade G. Regehr

**Author notes:** These authors contributed equally (Leannah N. Newman, Amanda P. Stark).

## Abstract

In addition to its well-known contributions to motor function, the cerebellum is involved in emotional regulation, anxiety, and affect^1-4^. We found that suppressing the firing of cerebellar Purkinje cells (PCs) rapidly excites forebrain areas that could contribute to such functions (including the amygdala, basal forebrain, and septum), but that the classic cerebellar outputs, the deep cerebellar nuclei (DCN), do not project to these forebrain regions. Here we show that PCs directly inhibit parabrachial nuclei (PBN) neurons that project to and influence numerous forebrain regions in a manner distinct from the DCN pathway. We also found that the PBN and DCN output pathways differentially influence behavior: suppressing the PC to PBN pathway is aversive and does not affect the speed of movement, whereas suppressing the PC to DCN pathway is not aversive and reduces speed. Molecular profiling shows that PCs inhibit numerous types of PBN neurons that control diverse nonmotor behaviors^5-9^. Therefore, the PBN pathway allows the cerebellum to regulate activity in many forebrain regions and may be an important substrate for nonmotor disorders related to cerebellar dysfunction.

---

The posterior vermis of the cerebellar cortex has been implicated in many nonmotor behaviors. In animal models, disruption or stimulation of different regions of the vermis modulates aggression^10^, motor planning^11,12^, spatial memory^13-15^, aspects of fear^16-18^, and hippocampal epilepsy^19,20^. In humans, damage to the vermis is associated with deficits in emotional control, language, memory, and executive function^3,4^. Cerebellar damage can also result in emotional disturbances consistent with limbic system dysfunction^1,2^.

It is not known how the posterior vermis influences these behaviors, but it seems likely that it somehow influences regions associated with them. Electrical stimulation of the cerebellum rapidly increases activity in the hypothalamus^21,22^, the amygdala^22-24^, basal forebrain^22^, septum^22,24^, hippocampus^22-25^, and cortical regions^21-23^. The interpretation of these results is complicated because electrical stimulation can antidromically activate mossy fibers and modulatory fibers.

To assess whether activity in posterior vermis of the cerebellum influences other brain regions, we used optogenetics to selectively suppress spontaneous PC firing. This selectively disinhibits PC targets and is more selective than electrical stimulation. We briefly (20 ms) illuminated the posterior vermis through a thinned skull to suppress PC firing in awake head-restrained mice that express halorhodopsin in PCs, and monitored activity in downstream regions with multielectrode arrays. Suppressing PC firing elevated spiking in 62% of thalamic neurons (4.2±1.0-fold increase, 28±1 ms latency) in regions known to receive DCN projections^26-28^ (**Fig. 1b, Extended Data Fig. 1aef**). PC suppression also increased spiking in 41% of amygdala neurons (2.8±0.7-fold increase, 33.4±1.8 ms latency, **Fig. 1b, Extended Data Fig. 1bef**), in 54% of neurons in the septum (3.0±0.5-fold increase, 33±2 ms latency, **Fig. 1b, Extended Data Fig. 1cef**). In the basal forebrain, 21% of cells increased firing (2.0±0.3-fold, 23.3±2.8 ms latency) and 31% of the cells were inhibited (20±2 ms, latency **Fig. 1b, Extended Data Fig. 1d-f**). These results establish that suppressing PC firing in the cerebellar vermis rapidly influences firing in the thalamus, the amygdala, the septum, and the basal forebrain. These findings motivated us to determine the output pathways that allows the vermis to influence such forebrain regions.

**Fig. 1:**
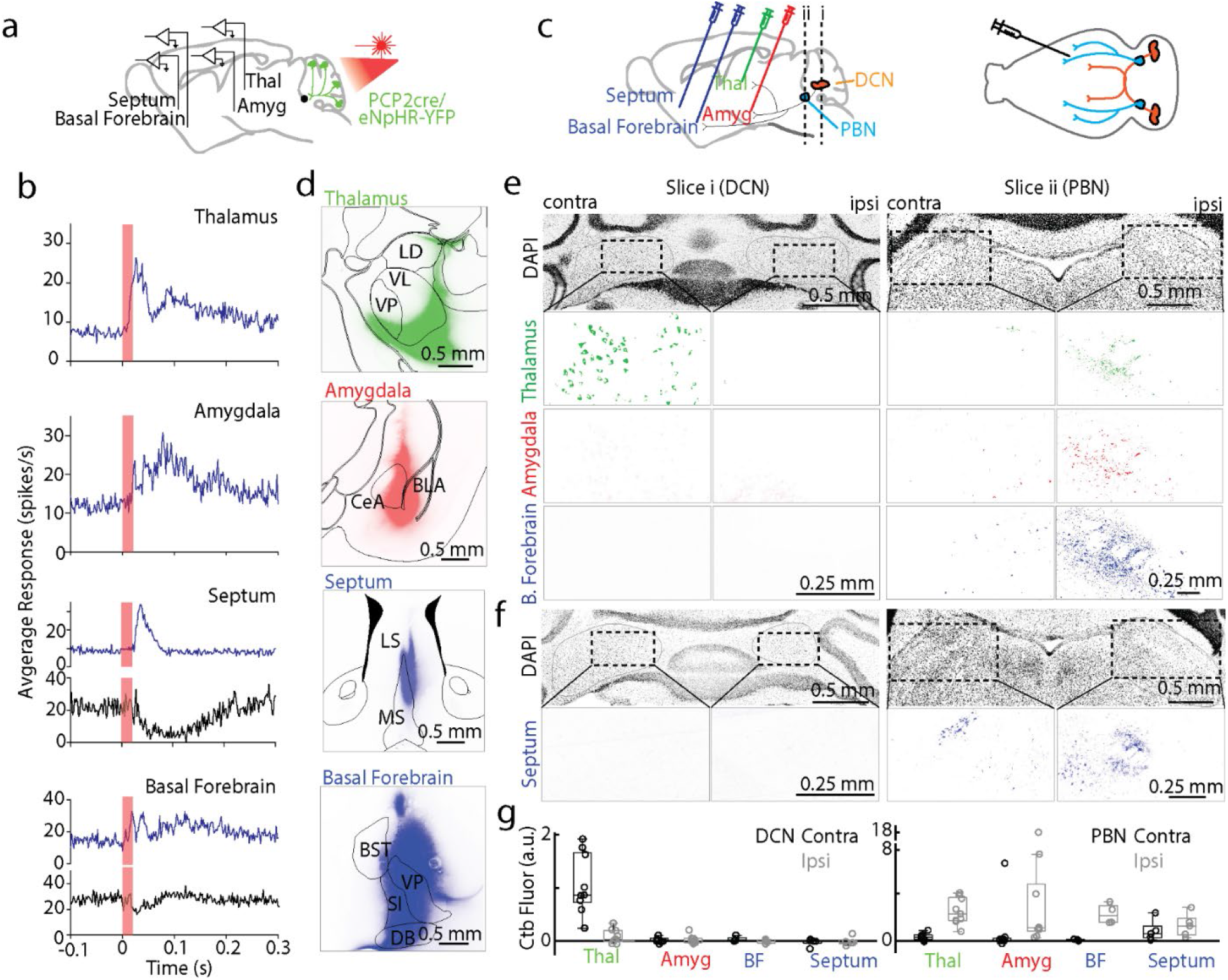
Suppressing Purkinje cell firing evoked short latency responses in multiple brain regions that are directly innervated by the PBN but not the DCN. **a**. Single-unit, multielectrode array recordings were made from awake, head-restrained PCP2Cre/Halo mice across several areas in the brain. 20 ms pulses of light were delivered to the posterior cerebellum. **b**. Average increases in activity from recordings in the thalamus (54/87 responding neurons), amygdala (30/73), septum (37/68), and basal forebrain (18/86) are shown in blue. Decreases in activity (black) were seen in the septum (4/68) and basal forebrain (27/86). **c**. (Left) Retrograde tracer cholera toxin subunit B (Ctb) labelled with different color fluorophores was unilaterally injected into the thalamus, amygdala, basal forebrain, or septum. (Right) The deep cerebellar nuclei (DCN) and parabrachial nuclei (PBN) were then examined for retrograde labelling. **d**. Injection sites are shown for thalamus, amygdala, and basal forebrain done in a single animal, and for the septum in a different animal **e**. Retrograde labelling in the DCN and PBN is shown for an animal with three injections sites in **d**. **f**. Retrograde labelling in the DCN and PBN for the animal with septum injection site in **d**. **g**. Quantification of retrograde labelling (Ctb fluorescence – background fluorescence) observed following injections into the indicated regions: thalamus injections (n=9), amygdala (n=9), basal forebrain (n=4), and septal injections (n=5). For all box plots: central mark of each box is the median, the edges represent the 25^th^ and 75^th^ percentiles, and the whiskers represent the range of data. LD: laterodorsal, VL: ventrolateral, VP: ventral posterior, CeA: central amygdala, BLA: basolateral amygdala, LS: Lateral septum, MS: medial septum, BST: bed nucleus of the stria terminalis, VP: ventral pallidum, SI: substantia innominate, and DB: diagonal band of broca.

To provide insight into the direct synaptic inputs to these forebrain regions of interest, we injected different color variants of cholera toxin subunit β, a retrograde tracer^29^, into the thalamus (n = 9), amygdala (n = 9), basal forebrain (n = 4), and septum (n = 5) (**Fig. 1c**). Several days later, we sliced sections to determine the injection site (**Fig. 1d**), and sectioned the cerebellum and brain stem to identify retrogradely labelled cells (**Fig. 1e-f**). Thalamic injections retrogradely labelled many neurons in the contralateral DCN, and few cells in the ipsilateral DCN, which is consistent with the well described direct DCN to thalamus pathway ^26-28^. In contrast, injections into the amygdala, basal forebrain and the septum did not show any retrograde labelling in the DCN (**Fig. 1e-g**), but led to prominent labelling of the ipsilateral PBN and little to no labelling in the contralateral PBN (**Fig. 1e, g**). The close proximity of the septum to the midline made it difficult to restrict injections to just one side of the septum to assess the extent of contralateral labelling. Thalamic injections also labelled the ipsilateral PBN (**Fig. 1e, g**).

The labelling of the PBN is intriguing because some PCs in the vermis directly synapse within the PBN^30-33^. However, it was thought that this projection primarily controls autonomic functions like heart rate and respiration through descending projections from the PBN^30-33^, and there was no indication that the PC to PBN pathway allows the cerebellum to influence the forebrain. However, regions of the PBN project to the forebrain^34^, and the PBN is implicated in behaviors related to cerebellar dysfunction^35,36^, so we hypothesized that the PC to PBN pathway influences the forebrain.

To determine the regions of the cerebellar cortex that project to the PBN, we injected retrobeads into the PBN, and found that they retrogradely labelled PCs primarily in the posterior lobules, VIII-X (**Fig. 2a-b**), in agreement with a previous study^30^. The highest density of labelled PCs was ipsilateral to the injection site 0.56 ± 0.06 mm from the midline (**Fig. 2c**).

**Fig. 2:**
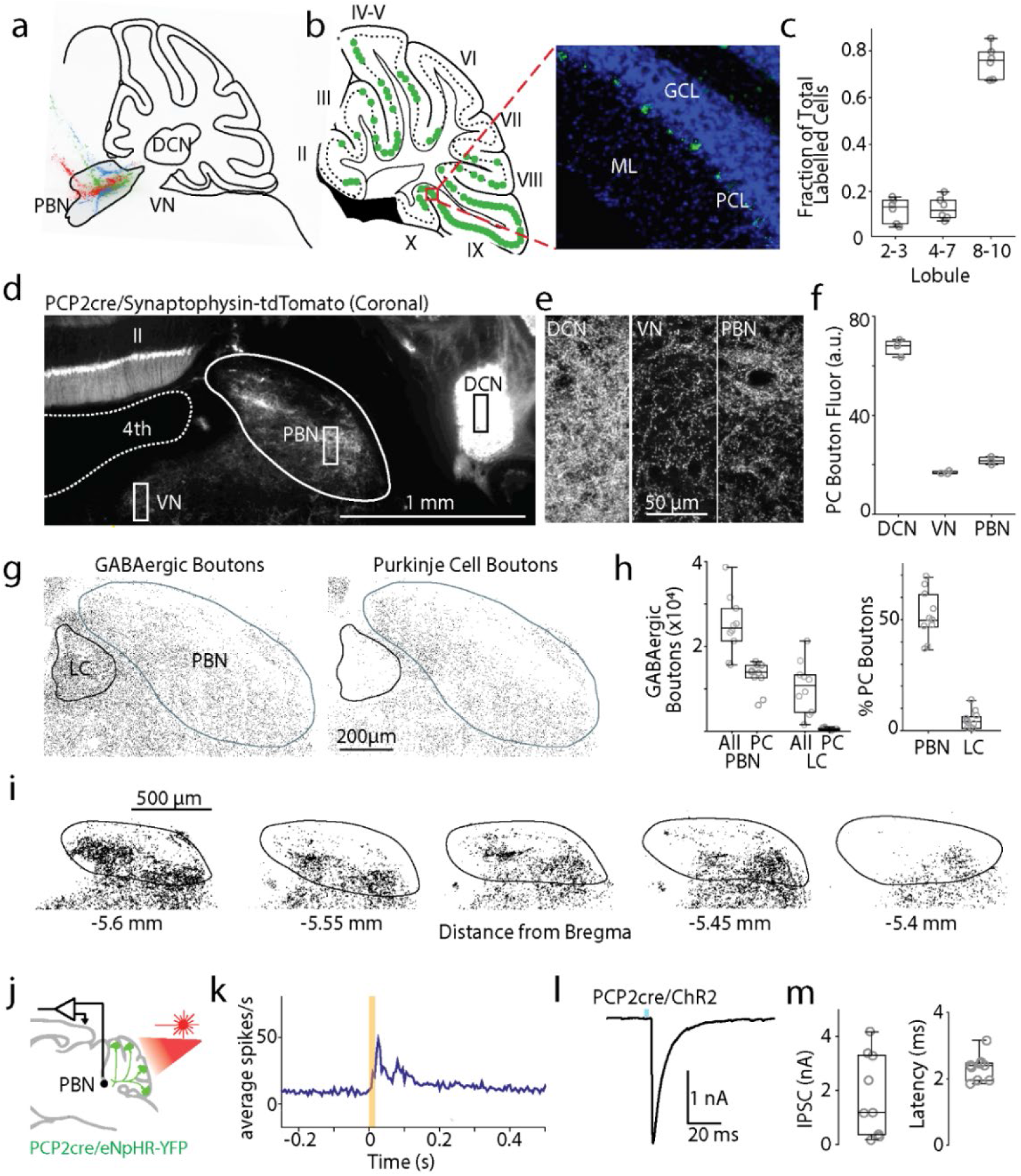
Purkinje cells make numerous synapses within the PBN. **a**. Fluorescent retrograde beads were injected into the PBN (n= 3 mice, a different color for each injection). **b**. *left*, Location of labelled PCs from one injection (*green*). Cerebellar lobules II-X are indicated. *right*, Image of retrobead fluorescence (*green*) and DAPI labelling (*blue*). GCL: granular layer; PCL: PC layer; ML: molecular layer. **c**. Locations of labelled PCs. **d**. PCP2-cre Synaptophysin-tdTomato mice were used to label PC presynaptic boutons (some somatic and dendritic labelling was also present). 4th: 4th ventricle. **e**. Fluorescence images of regions as indicated in **d**. **f**. Quantification of PC bouton fluorescence in different regions. **g**. Left: staining for all GABAergic boutons (VGAT) around the PBN and LC. Right: GABAergic boutons overlapping with PCP2cre/tdTomato axons. **h**. Quantification of all GABAergic boutons in PBN and LC, and % of GABAergic boutons from PCs. **i**. Identified PCP2-Cre Synaptophysin-tdTomato boutons are shown for a series of slices. **j**. Single-unit, multielectrode array recordings were made in the PBN and the DCN in awake, head-restrained PCP2cre/Halo mice. The posterior cerebellar cortex was optically stimulated through a thinned skull (20 ms). **k**. Average firing evoked in rapidly responding PBN neurons (13/28). **l**. Optically-evoked synaptic current (1 ms blue light) in a PBN neuron recorded in brain slice from a PCP2-Cre ChR2 mouse. **m**. Summary of the amplitudes and latencies of light-evoked PC to PBN neuron synaptic current

To determine the extent of PC synapses in the PBN, we used PCP2-cre/Synaptophysin-tdTomato mice to fluorescently label all PC presynaptic boutons ^37,38^. Synaptophysin-tdTomato fluorescence was apparent in the DCN, the vestibular nuclei (VN), and in the PBN. (**Fig. 2d-f, Extended Data Fig. 2a**). The density of PC boutons in the PBN is comparable to the VN, and lower than in the DCN (**Fig. 2e, f**). We also compared the densities of PC synapses in the PBN and in the nearby LC, another PC target^39^. We used TH labelling to delineate the LC, vGAT immunofluorescence to identify all GABAergic synapses, and labelled PC axons genetically (**Fig. 2gh, Extended Data Fig. 2c**)^40^. Inhibitory synapses are widespread in the LC and PBN, but there are approximately 40 times more PC synapses in the PBN than in the LC, and a much higher fraction of these synapses are from PCs in the PBN than in the LC (**Fig. 2h**, *right*). These suggest that PCs have a much stronger influence on the PBN than on the LC. We found that PC synapses are present at higher densities on the medial side of the brachium conjunctivum in the coronal plane, and in posterior regions (**Fig. 2i, Extended Data Fig. 2b**). The widespread but heterogeneous distribution of PC synapses within the PBN suggests that the cerebellum regulates many, but not all, PBN-dependent behaviors.

We used optogenetics to test the properties of PC inhibition of PBN neurons both *in vivo* and *in vitro*. We suppressed PC firing *in vivo* in PCP-cre/halorhodopsin mice (**Fig. 2j, Extended Data Fig. 3ab**). and found that firing was elevated in 68% of PBN neurons, with a short latency in 47% of cells (**Fig. 2k, Extended Data Fig. 3c, f**), and with a long latency in 21% (**Extended Data Fig. 3d, f**). In similar experiments for DCN neurons, firing was evoked in all DCN neurons with a short-latency (<30 ms) (**Extended Data Fig. 3e, f**). There was no secondary elevation of activity within the DCN, where recurrent excitation has not been described. These results suggest that in contrast to the DCN where PCs inhibit all cells, PCs only inhibit about half of the cells in the PBN. The long-latency responses evoked in 21% of PBN neurons suggest that these neurons are not directly inhibited by PCs, but they may be disynaptically excited. To more definitively determine whether PBN neurons are directly inhibited by PCs, we characterized the PC to PBN synapse in acute coronal PBN brain slices using PCP2-cre/ChR2 mice while blocking excitatory synaptic transmission. Optical stimulation of PCs evoked large (1.8±0.5 nA), short latency (2.33±0.13 ms) IPSCs in 9 of 12 PBN neurons (**Fig. 2l-m**), indicating that PCs powerfully and directly inhibit approximately 75% of the neurons in the posterior PBN.

To assess the functional roles of the PC to PBN pathway the PC to PBN projection must be targeted selectively. Suppression of the posterior vermis is not sufficiently selective because it also affects projections to the DCN and the VN. However, we found that inhibiting halorhodopsin-expressing PC axons with an optrode evoked rapid (5.8±0.9 ms latency), increases (1.5±0.1 fold) in 54% (17/31) of PBN neurons (**Fig. 3ab, Extended Data Fig. 4**). We used the immediate early gene c-Fos as an activity reporter ^41,42^ to assess the spatial extent of stimulation. We unilaterally illuminated PC axons in either the PBN (n=4) or the DCN (n=3) (100 ms every 8 s for three hours) and then stained for c-Fos (**Fig. 3c-d**). PC suppression elevated c-Fos near the tip of optical fibers in either the PBN (**Fig. 3c**) or the DCN (**Fig. 3d**). PC-PBN suppression did not elevate c-Fos staining relative to control animals (no halorhodopsin) in the contralateral PBN, the LC (a region with high rates of spontaneous firing), the DCN, or the VN (**Fig. 3ef, Extended Data Fig. 5**). DCN stimulation elevated c-Fos staining in the ipsilateral and contralateral DCN, but did not elevate c-Fos staining in either the PBN or the VN (**Fig. 3ef, Extended Data Fig. 5**). These findings suggest that we can use this approach to selectively suppress specific pathways, but that it only excites the small fraction of cells near the tip of the optical fiber.

**Fig. 3:**
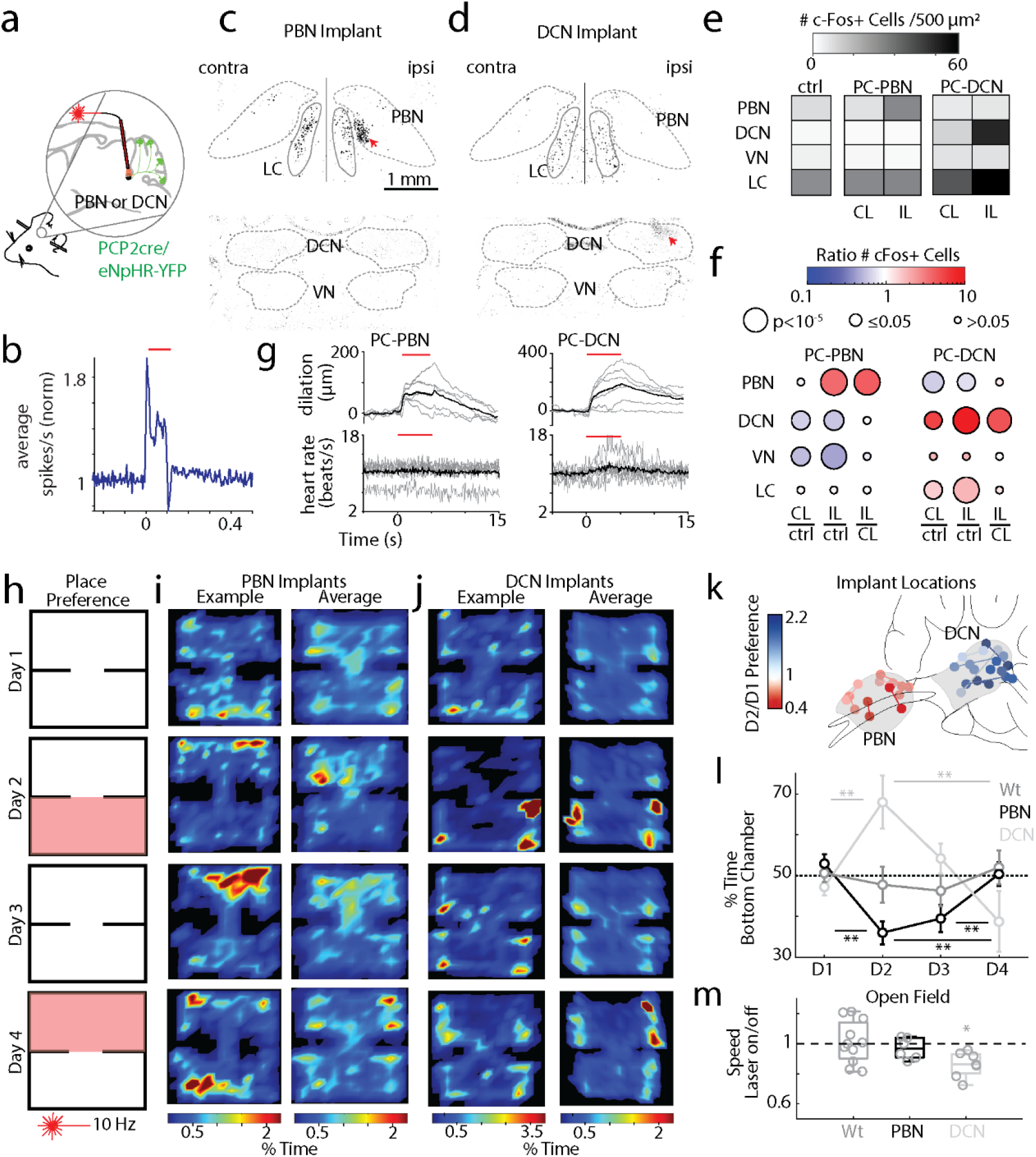
Suppression of the PC-PBN pathway is aversive. **a**. Schematic of Halo/PCP2cre mice with bilaterally implanted optical fibers in either the PBN or DCN. **b**. Average single-unit PBN response to the inhibition of the PC-PBN pathway (red line) (n=17). **c-d**. Suppressing PC-PBN or PC-DCN pathways locally elevated c-Fos expression (red arrows). **e**. Summary of number of c-Fos expressing cells in different regions after PC-PBN, PC-DCN, or control (wildtype) stimulation. CL: contralateral to stimulation; IL: ipsilateral. **f**. Comparisons of c-Fos expression shown in **e**. Cool colors indicate a decrease in expression and hot colors an increase. Statistical significance for each comparison is indicated by symbol size (**Table 1**). **g**. The effects of suppressing either the PC-PBN or PC-DCN pathways on pupil dilation and heart rate (**Table 2**). **h**. Experimental configuration is shown for a two-chamber place preference test. Regions in which PC inputs to either the DCN or the PBN were optically stimulated are indicated in red. **i**. Example (left) and average (right, n=7) position heat maps for corresponding test days for PC-PBN suppression. **j**. Same as **i**, but for PC-DCN suppression (n=9). **k**. Bilateral implant locations. Chamber preference for each animal (Test/Baseline; D2/D1) is encoded by the color of the corresponding implant location pairs. **l**. Summary of the % time spent in the bottom chamber for PC-PBN suppression (red) and DCN suppression (blue), and control mice (grey) (mean ± S.E.M, **Table 3**). **m**. Change in velocity after stimulation in open field test (see **Extended Data Fig. 7, Table 4**).

We compared the behavioral effects of the PC-PBN and PC-DCN pathways. We began with the control of the autonomic system, which has been thought to be the primary function of the PC-PBN pathway ^31,43-45^. We found that bilaterally suppressing either the PC-PBN or the PC-DCN pathway (5s, 10 Hz, 50 ms illumination, **Extended Data Fig. 6, Table 2**) dilated pupils (**Fig. 3g**, *top*). Suppressing the PC-DCN pathway slightly elevated heart rate, but suppressing the PC-PBN pathway did not (**Fig. 3g**, *bottom*, **Table 2**). We then examined the effects of PC-PBN and PC-DCN stimulation in a place preference test (**Fig. 3h-l**). We hypothesized that these pathways might have very different effects because PBN activation is typically aversive ^46,47^, whereas DCN outputs can be rewarding ^48^. On Day 1, mice were placed in a two-chamber arena for 15 minutes. On Day 2, a pathway was optically suppressed (**Extended Data Fig. 6**) when mice were in the lower chamber. On Day 3, there was no stimulation and on Day 4, the pathway was optically suppressed when the mice entered the top chamber. Suppressing the PC-PBN pathway was aversive (fraction of time in lower chamber 0.36 ± 0.02. on Day 2 vs. 0.53 ± 0.02 on Day 1), preferences were maintained on Day 3 (0.40 ± 0.03), but were eliminated during the reversal trial on Day 4 (0.50 ± 0.03) (**Fig. 3 i, l, m, Table 3**). In contrast, bilateral suppression of the PC-DCN pathway increased the fraction of time in lower chamber (**Fig. 3j-l**, 0.68 ± 0.06 on Day 2 vs. 0.47 ± 0.02 on Day 1, **Table 3**). Wildtype control animals never showed a preference (**Fig. 3l**). To interpret these experiments, we used an open field assay to determine if suppression of either pathway influences speed (**Fig. 3m, Extended Data Fig. 7, Table 4**). Suppressing the PC-PBN pathway did not alter speed, but suppressing the PC-DCN pathway reduced speed by ∼15% (5.1 ± 0.6 to 4.4 ± 0.6 cm/s). Taken together, our findings indicate that suppressing the PC-PBN pathway is aversive, that there is a memory of this aversion, and that this memory can be reversed. This suggests that ongoing PC firing normally suppresses the firing of aversive PBN neurons. Suppression of the PC-DCN pathway has very different effects. The increased presence in the lower chamber could arise either from this pathway being rewarding ^48^, or from reduced speed.

We took two approaches to provide insight into how the PC-PBN pathway could influence the forebrain and affect behavior (**Fig. 4**). First, we used an anatomical transynaptic approach to label PBN neurons that receive direct PC inhibition. We injected AAV1-syn-Cre into the posterior vermis. This virus effectively expresses cre in anterograde targets, but also retrogradely labels cells with low efficiency ^49^. In the PBN, cre expression is restricted to anterograde targets because the PBN does not project to the vermis ^7,50^. We also injected AAV-flex-tdTomato in the PBN to label PBN neurons that are directly inhibited by PCs (**Fig. 4a**). tdTomato expressing fibers were found in the amygdala, basal forebrain, the septum and other forebrain regions (**Fig. 4b-d, Extended Data Fig. 9**), indicating that PC-recipient PBN neurons project directly to these regions.

**Fig. 4:**
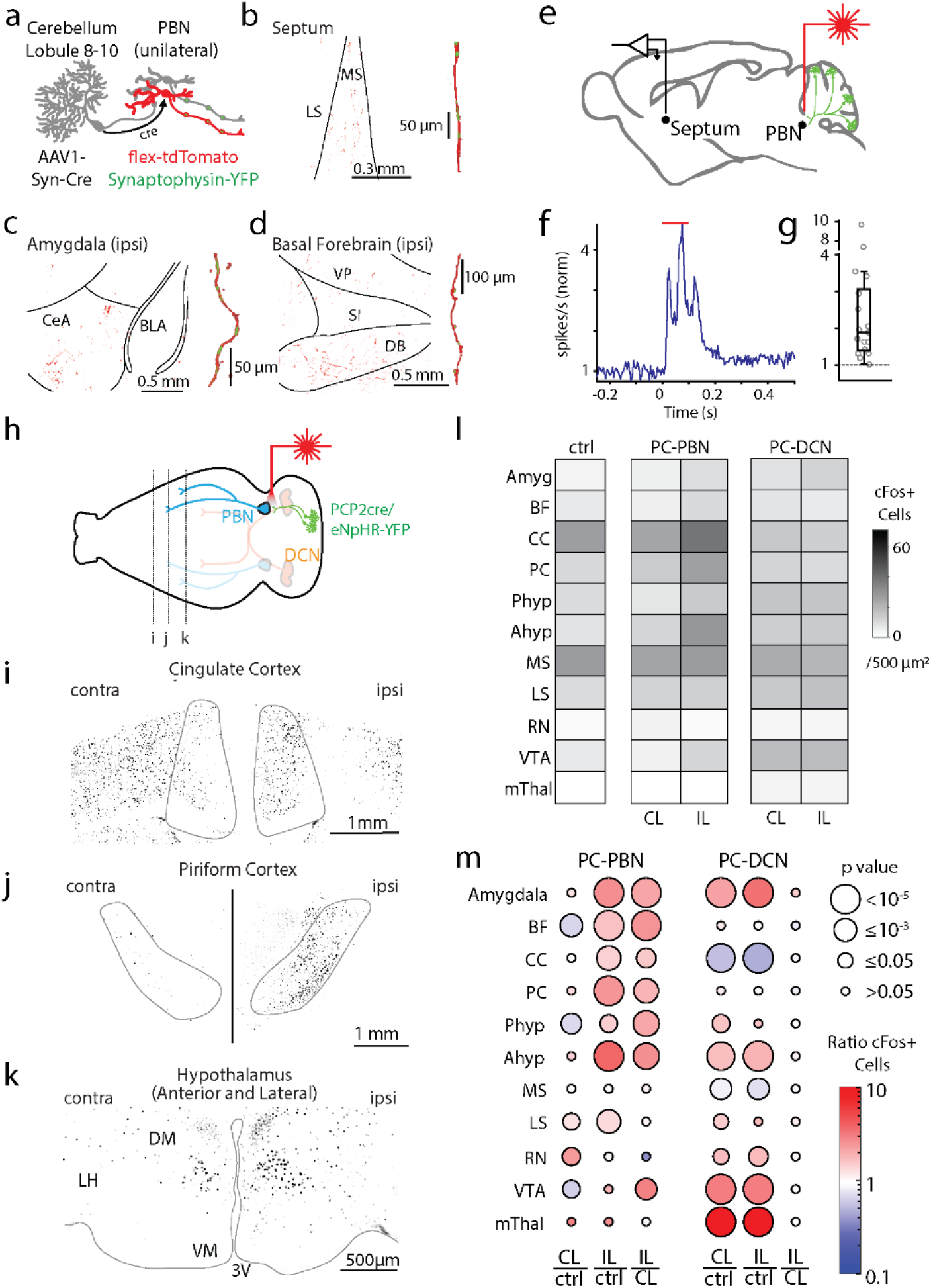
Purkinje cell-recipient PBN neurons project to and influence numerous forebrain brain regions. **a**. Injections of anterograde AAV-cre into the posterior vermis and AAVs with flex-tdTomato and synaptophysin-YFP and into the PBN, labelled PC-recipient PBN neurons with tdT and the presynaptic boutons of all PBN neurons with YFP. **b**. *left*, Low magnification fluorescence image of the septum. *right*, high magnification image of a reconstructed axon (red) in the septum and colocalized synaptophysin-YFP (green). **c**. Same as **b**, but for the amygdala. **d**. Same as **c**, but for the basal forebrain. **e**. Optical suppression (100 ms in Halo/PCP2-Cre mice) of the PC-PBN pathway elevates firing in the septum. **f**. Average response (n=17 cells). **g**. Distribution of normalized responses due to optical suppression of the PC-PBN pathway **h**. Schematic. Following unilateral optical suppression of either the PC-PBN or PC-DCN pathway, slices were stained for cFos. **i-k**. cFos staining for the coronal slices indicated in **h**. **l**. Quantification of c-Fos expression after ipsilateral (IL) PC-PBN, PC-DCN, or control (wildtype) stimulation. **m**. The ratio of c-Fos expression in the stimulated (IL) and unstimulated (CL) hemisphere was determined relative to control (ctrl) mice. The ratio IL to CL expression was also determined. The average ratio (color) and the statistical significance (symbol size) are indicated (**Table 1**). BF: Basal forebrain, CC: Cingulate Cortex, PC: Piriform Cortex, Phyp: Hypothalamus Preoptic Area. Ahyp: Hypothalamus Anterior and Lateral (as in **k**), MS: Medial Septum, LS: Lateral Septum, RN: Red Nucleus, VTA: Ventral Tegmental Area, mThal: Motor thalamus.

We also assessed the effect of the cerebellum-PBN pathway on downstream regions by selectively suppressing PC-PBN synapses and examining activity in downstream regions (**Fig. 4 e-m**). PC-PBN suppression evoked large (2.7±0.6-fold) short latency (21.6±2.1 ms) increases in firing in 17/40 septum neurons (**Figure 4e-g, Extended Data Fig. 10**). We also used cFos expression to determine the regions activated by suppression of either the PC-PBN pathway or the PC-DCN pathway (**Fig. 4 h-m**). Unilateral suppression of these pathways affected cFos expression very differently. Suppressing the PC-PBN pathway elevated cFos labelling in many ipsilateral forebrain regions (**Fig. 4h-m**), whereas suppressing the PC-DCN pathway evoked bilateral increases (**Fig. 4m, Extended Data Fig. 11, Table 1**). These pathways also affected different regions. Only PC-PBN suppression increased cFos expression in the basal forebrain, cingulate cortex, piriform cortex, and the preoptic area of the hypothalamus, whereas only PC-DCN suppression led to prominent increases in the motor thalamus and the VTA. Suppression of either pathway evoked increases in the amygdala, the anterior hypothalamus and the red nucleus (which is consistent with known direct and indirect DCN pathways ^48,51,52^). These results suggest that although there is some overlap, the PC-PBN pathways and PC-DCN pathways differentially influence various forebrain regions.

There are many types of neurons within the PBN that project to different regions and differentially influence behavior (**Table 6**). To identify the PBN neurons targeted by PCs within the PBN, we injected AAV1-Syn-Cre and AAV-CAG-tdT in the posterior vermis of Sun1-GFP mice to label PCs and their axons with tdTomato and to trans-synaptically label the nuclei of PC-targeted neurons with GFP (**Fig. 5a**). AAV1-Syn-Cre provide an effective means of anterogradely labelling target cells in the PBN, which does not project to the vermis ^7,50^. We observed many labelled nuclei in the PBN in the vicinity of PC fibers (**Fig. 5e**). Labeled nuclei were also present outside the PBN, far from labelled PC fibers. Because this approach can retrogradely label cells at low efficiency, regions that project to the cerebellar cortex such as the LC and VN ^7,39,50,53-55^ can be labelled. We tested whether PBN neurons that are inhibited by PCs in the vermis in turn project to the forebrain by injecting AAV1-Syn-Cre viruses in the cerebellar vermis of Ai75 mice to anterogradely label the nuclei of directly targeted PBN neurons (*red*), with injections of cholera toxin into the amygdala to retrogradely label cells (*blue*) (**Fig. 5f**). Numerous PBN neurons were colabelled, establishing that PCs directly inhibit PBN neurons that in turn project to the forebrain (**Fig. 5g**).

**Fig. 5.**
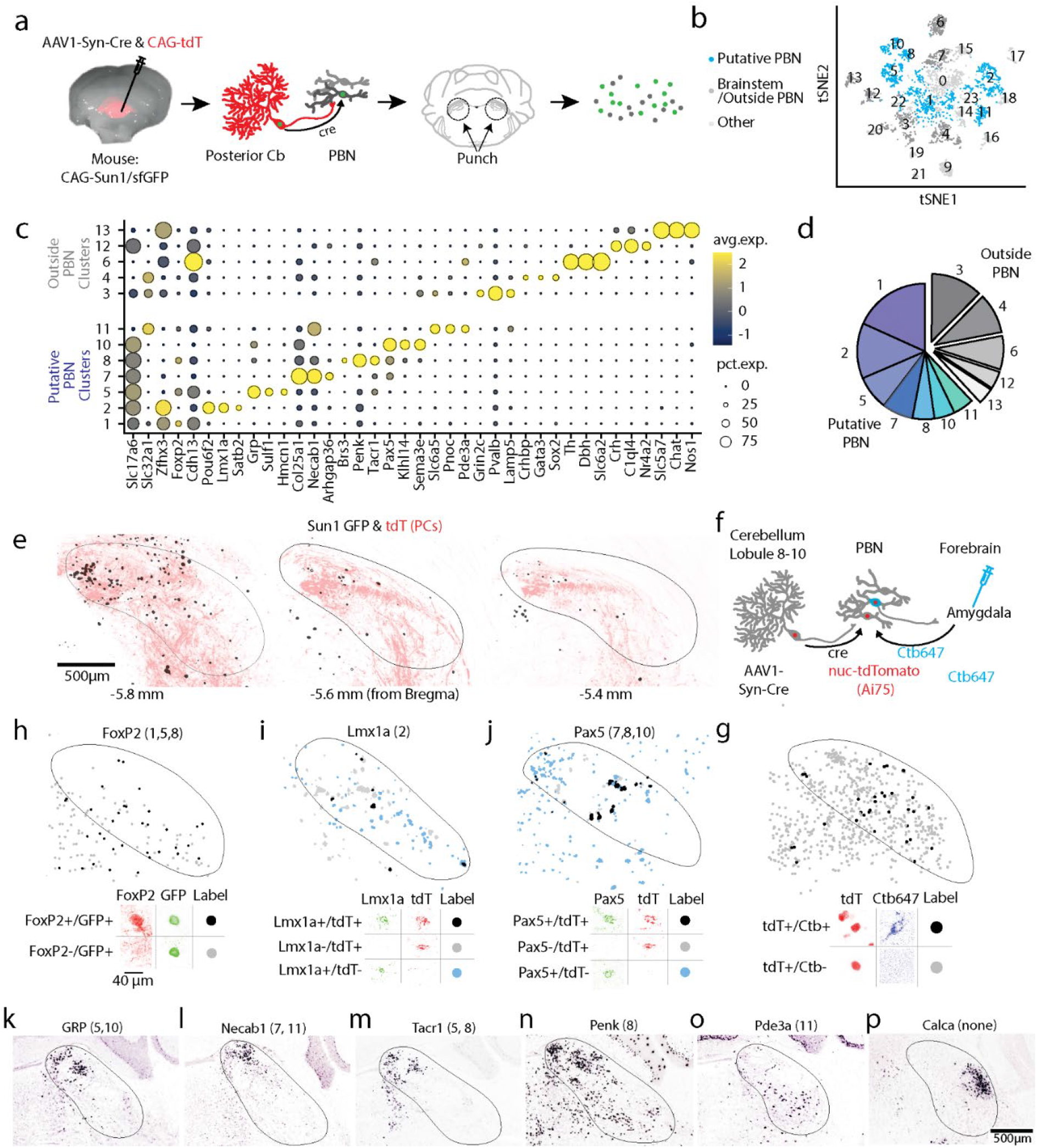
Diversity of PBN neurons directly targeted by PCs. **a**. Schematic showing the strategy used to label PC targets. AAV1-Syn-Cre was injected into the posterior vermis of a CAG-Sun1/sfGFP mouse. The nuclei of target PBN neurons were trans-synaptically labelled and sorted. **b**. tSNE visualization of 3876 neuronal nuclei separated into 23 neuronal clusters (**Extended Data Fig. 12)** **c**. Dot plot of scaled expression for indicated clusters. Clusters with cerebellar markers, unclustered nuclei, and clusters of < 100 nuclei were removed (**Extended Data Fig. 12)** **d**. Fraction of labelled nuclei per cluster. **e**. Labelling of nuclei (black) and PC axons (red) in 3 coronal planes following an injection as in **a**, along with a AAV1-CAG-tdTomato coinjection. **f**. Schematic showing the strategy used to retrogradely label cells projecting to the amygdala and anterogradely label PBN neurons directly inhibited by PCs. **g**. (*top*) A fraction of TdT + neurons were retrogradely labelled by cholera toxin injected into the amygdala (*black*) and many were not (*grey*). (*bottom*) Representative cells showing labelling and the corresponding symbols used to present cell locations. **h**. (*top*) A fraction of trans-synaptically labelled PBN neurons were Foxp2+ (*black*) and many were Foxp2-(*grey*). (*bottom*) Representative cells showing Foxp2 immunohistochemistry and nuclear labeling. **i**. *In situ* hybridization for Lmx1a and tdTomato labelled a fraction of the trans-synaptically labelled cells (black) in a tdTomato reporter mouse (Ai75d). Labelled cells were often found in groups of 3-4 cells. **j**. Same as (**g**) but for Pax5. **k-p**. Images from the Allen Brain Atlas staining for selected genes corresponding to indicated clusters. Calca labels CGRP neurons in the PBN that were not trans-synaptically labelled.

To gain insight into the molecular properties of PC-targeted PBN neurons, we injected AAV1-Syn-Cre in Sun1-GFP mice, microdissected the PBN and nearby tissue, isolated GFP-labelled nuclei, and used single-nucleus RNA-seq (snRNAseq) to profile their transcriptome (**Fig. 5a**). Of 3876 neurons, 3300 neurons were grouped into 23 clusters characterized by uniquely expressed genes (i.e., markers) (**Fig. 5b, Extended Data Fig. 12, Table 5, Supplementary Data 1**), whereas the remaining ones did not have a clear marker (cluster 0). We identified markers associated with clusters comprised of over 100 neurons (**Fig. 5c**), and observed trans-synaptic labelling of neurons in the PBN that expressed FoxP2 (clusters 1, 5, 8, **Fig. 5f**), Lmx1a (cluster 2, **Fig. 5g**), or Pax5 (clusters 7, 8, 10, **Fig. 5h**). Based on these experiments, and *in situ* hybridizations from the Allen Brain Atlas^56^ (**Fig. 5i-n, Table 5**), we conclude that 7/12 clusters were likely PBN neurons and the remaining were comprised of neuron types outside the PBN (**Fig. 5d**). Although PCs directly inhibit many types of PBN neurons, the absence of CGRP (*Calca* gene) neurons ^57^ in our clusters (**Fig. 5p, Supplementary Data 1**) suggests that PCs may not inhibit all types of PBN cells.

Identifying the PBN cell types targeted by PCs, allowed us to link our findings to recent studies in which Cre driver lines and localized injection of AAVs in the PBN were used to determine the projections of PBN neuron subtypes (**Table 6**). Glutamatergic (*Slc17a6*) PBN neurons (comprising all PBN projection neurons and present in clusters 1, 2, 5, 7, 8 and 10) have widespread projections to the cerebral cortex, the septum, basal ganglia, amygdala, the thalamus and the midbrain ^7^. *Penk* PBN neurons (clusters 1, 2, 5 and 8) project to the hypothalamus ^9^ and *Foxp2* PBN neurons (clusters 1, 5, 8, 11) project to the septum, basal ganglia, amygdala, basal forebrain, thalamus, and hypothalamus^7^. *Slc17a6* and *Penk* PBN neurons are both aversive in a place preference assay^9^, consistent with our observation that PC-PBN pathway suppression is aversive (**Fig. 3i,k-m**). PBN *Satb2* neurons (cluster 2) respond to different tastes, and optogenetic activation of PBN *Sat2b* neurons enhances taste preferences ^8^. *Tacr1* PBN neurons (clusters 5, 7 and 8) regulate the response to ongoing pain and itch ^5^. As more is learned about how specific types of PBN neurons regulate behaviors, our findings will provide insight into how these behaviors are regulated by the PC-PBN pathway.

Our main finding is that the cerebellum has a much more extensive and powerful influence on the PBN than had been previously appreciated, and that the PC to PBN output pathway allows the cerebellum to influence many forebrain regions that are involved in diverse nonmotor behaviors. Optogenetic suppression of PC spiking in the posterior vermis powerfully disinhibits more than half of posterior PBN neurons, and causes them to spike rapidly. This suggests that PCs, which fire spontaneously *in vivo* from 20-120 Hz, provide ongoing suppression of spiking within the PBN. We used numerous experimental approaches to overcome the many challenges involved in studying the PC-PBN pathway (**Table** 7).

One of the most distinctive features of the cerebellar output pathway through the PBN is that it projects extensively throughout the forebrain, whereas the conventional PC to DCN pathway does not. We established this through trans-synaptic anterograde tracing, c-Fos, and *in vivo* electrophysiology. These targets include many brain regions not normally associated with cerebellar function, motor performance or motor learning, such as the amygdala, basal forebrain, hypothalamus and the septum. The PC-PBN pathway provides a long sought-after explanation for the many studies suggesting that the cerebellum plays an important role in emotional regulation, possibly through a direct cerebellar pathway to limbic regions ^21-25^. Our place-preference experiments indicate that disinhibition of the PC-PBN pathway is aversive whereas disinhibition of the PC-DCN pathway is not. This allows simple pauses in PC firing full control over the entire spectrum of valence, where previous work focused on their role in reward ^48^.

A number of additional cerebellar pathways exist that allow it to modulate valence. Recent work ^17,18^ has shown that bidirectional modulation of DCN activity can bidirectionally modulate aspects of fear. Increases in DCN activity lead to profound fear extinction and decreases prolong fear memories, in line with the valence of outputs indicated in this study. Several other output pathways from the DCN have been proposed (to hypothalamus^51^, substantia nigra, and others), though their function has yet to be determined.

Damage to the posterior cerebellum causes symptoms consistent with the function of PC-PBN projection targets. Cerebellar Cognitive Affective Syndrome (CCAS) is a syndrome resulting from damage (genetic or otherwise) to the posterior lobules of the cerebellum^3^. These symptoms include disturbances in affect, emotional control, arousal, and executive function, and correspond very well with known roles of the regions targeted by PC-PBN outputs that we describe here. The basal forebrain plays an important role in affect, the amygdala is critical for emotional control, activity in the septum correlates with arousal, and the cingulate cortex is a part of the prefrontal, executive cortices. In contrast, the DCN only have polysynaptic inputs to these regions. Thus, the PC-PBN pathway is well suited to playing an essential role in behaviors associated with CCAS. The posterior cerebellum and the PC-PBN pathway is also particularly sensitive to concussive injury^58,59^, in particular the cerebellar lobules lining the 4^th^ ventricle. Traumatic brain injury can also damage the tracts of the superior cerebellar peduncle, where the cerebellar axons course through and the cells of the PBN reside. This type of damage is most commonly associated with PTSD and anxiety disorders^60^. Both of these conditions are tightly linked to the downstream structures in the PC-PBN pathway. In addition to CCAS and PTSD, clinicians have more broadly observed cerebellar associations with sleep disturbances^61^, anxiety^62^, schizophrenia^63-65^, and mood disorders^66^. The PC-PBN pathway provides multiple avenues for the cerebellum to influence the limbic system and associated neurological disorders.

## Materials & Methods

### Animals used

For injection of retrograde and anterograde tracers (cholera toxin, retrobeads, and AAVs), male and female C57BL/6J mice (Jackson labs) were used. For in vivo electrophysiology and behavior, male and female B6.Cg-Tg(Pcp2-cre)3555Jdhu/J x B6;129S-Gt(ROSA)26Sor^tm39(CAG-hop/EYFP)Hze^/J (PCP2-cre x Halorhodopsin, Jackson Labs) were used. For the anatomical tracing studies, male and female *B6*.*Cg-* Tg(Pcp2-cre)3555Jdhu/J x B6;129S-Gt(ROSA)26Sor^tm34.1(CAG-Syp/tdTomato)Hze^/J (PCP2-cre x synaptophysin-tdTomato, Jackson Labs) or B6.Cg-Tg(Pcp2-cre)3555Jdhu/J x B6.Cg-Gt(ROSA)26Sor^tm9(CAG-tdTomato)Hze/J^ (PCP2-cre x tdTomato, Jackson Labs) were used. For isolation of nuclei for RNA sequencing, male and female B6;129-Gt(ROSA)26Sor^tm5(CAG-Sun1/sfGFP)Nat/J^ mice were used (Sun1-GFP, Jackson Labs). For tdTomato labelling of nuclei in combination with in situ hybridization experiments and cholera toxin, B6.Cg-Gt(ROSA)^26Sortm75.1(CAG-tdTomato*)Hze/J^ were used (Ai75D, Jackson Labs). All animals were used under supervision of Harvard Medical School’s Institutional Animal Care and Use Committee (IACUC). We used the Mouse Brain in Stereotaxic Conditions Atlas^67^ as a reference for surgical coordinates.

### General Surgery Protocol

Mice were anesthetized and maintained under 2% isoflurane. Mice were secured to a Stereotaxic Surgery Instrument (Model 940 Small Animal, Kopf Instruments, Tujunga, CA). Eye ointment was applied and reapplied throughout surgery as needed. The heads of the mice were sanitized with an alcohol-coated wipe, after which the hair on the surgical area was removed with Nair. The exposed skin of the head was then sanitized with betadine solution. An incision was made to expose the cranium, and the surgery would proceed with the injection or implantation. Incisions were closed with sutures or dental cement (MetaBond, Parkell). Mice were injected with slow-release buprenorphine for analgesia and monitored for the next three days for post-op care.

### Preparation of mice for in vivo recordings and suppression of PC firing

For experiments in which PC firing in the posterior vermis was optogenetically suppressed (**Fig. 1a-b, Fig. 2jk, Extended Data Figs. 1, 3**), mice were implanted with a custom-made titanium head bracket and the cranium above the cerebellum and the recording areas of interest was exposed. During surgery, the skull above the cerebellum was thinned using a handheld drill until the underlying brain region was visible. After every pass with the drill, ACSF was dripped onto the skull to avoid thermal damage to surface neurons. At the end of the surgery, all areas exposed were covered with silicone elastomer (Kwik-Sil, World Precision Instruments). Mice were allowed to recover from surgery for at least three days. Mice were head restrained over a free-moving wheel for 30 minutes every day for 3 days prior to the first day of recording.

This preparation minimally perturbed cerebellar tissue and enabled us to stimulate a large area (approximately 3 mm in diameter) over multiple recording sessions, and has several advantages over electrical stimulation, which has been used in previous studies. It takes advantage of the fact that PCs fire spontaneously at high frequency, and suppressing this activity leads to rapid disinhibition that evokes rapid increases in firing of downstream neurons. In addition, because halorhodopsin is restricted to PCs, it specifically manipulates just those neurons, and not fibers of passage as in electrical stimulation. However, estimating the precise penetration of light through tissue in these experiments is challenging, as the cerebellum itself is heterogeneous in density, the thinning is variable, and the scattering is strongly affected by any vasculature in the region. Our rough estimate, based on Al-Juboori et al., 2013^68^ and Yizhar et al., 2011^69^, is that the light should be effective at most up to 2 mm past the surface of the skull, corresponding to most of the posterior cerebellum (lobules 6-9). This approach allowed us to assesses whether the firing PCs in the cerebellar vermis regulates firing in the amygdala, the septum and the basal forebrain similarly to the manner in which they influence firing in the thalamus. These experiments motivate the rest of the study, but they do not provide insight into the pathway or the complexity of that pathway, that allows the vermis to regulate activity in these regions.

### In vivo electrophysiology

Mice were anesthetized with 2% isoflurane. We then drilled a craniotomy over the recording site to exposed the desired brain area. After allowing the mouse to wake and recover for at least two hours, single-unit, multielectrode recordings were made with a silicon probe (P or E-style 16 channel probes, Cambridge NeuroTech) dipped in Di-I (Vybrant Multicolour Cell Labelling Kit, Thermofisher) while the mouse was head restrained over a freely rotating wheel. This procedure was repeated for a maximum of three days of recording per mouse. Once recordings were complete, mice were perfused with PBS and 4% PFA, 100 μm coronal slices were made from the brain tissue to determine electrode placement. Optrodes used in **Fig. 3b** were either constructed by gluing a 100 μm optical fiber (0.22 NA, Thorlabs) to a silicon probe (P or E-style 16 channel probes, Cambridge Neurotech) or custom-ordered from Cambridge Neurotech (P-style 16 channel probe with a Lambda-B tapered optical fiber attached).

### Optical stimulation

An MRL-III-635L Diode Red 635 nm Laser (Opto Engine LLC, Midvale, UT) was used to activate halorhodopsin. For experiments involving inhibiting PC soma through the thinned cranium, the beam was widened to encompass the entire thinned region that included the posterior vermis and paravermis. Steady-state power density from the laser was ∼80 mW/mm^2^. For experiments involving implanted optical fibers or optrodes, the steady-state intensity of light was ∼25 mW, as measured from the tip of the optical fiber prior to implantation or insertion.

### In Vivo Recording & Analysis

Data were sampled at 20 kHz using an RHD2000 recording system (Intan Technologies), and and bandpass filtered (0.1 Hz - 8 kHz). Data were sorted using Plexon Offline Sorter (Plexon Inc, Texas). Further analysis was all done in MATLAB (Mathworks, MA). Peri-stimulus firing rate histograms were generated from these spike times. Excitation or inhibition was defined as an increase or decrease of the firing rate (6 ms window) 2 standard deviations from the baseline firing rate after stimulus onset.

### Viral Injections

A hole was drilled at the desired medial-lateral (ML) and anterior-posterior (AP) coordinates for injection (Nanoject III, Drummond Scientific). The glass micropipette was placed 100 μm below the dorsal-ventral (DV) coordinate to create a “pocket” for the substance to be distributed. To minimize labelling of the injection tract, injections proceeded slowly (< 3 nl/s), and the pipette was left in place for at least 5 minutes following injection. Afterwards, the pipette was raised 100 μm and set in place for another 5 minutes to ensure that the substance was deposited at the injection site. The micropipette was then slowly retracted from the brain tissue.

### Choleratoxin injections

Wildtype mice were injected with 200 nL cholera toxin subunit B (CTB) in the thalamus (CTB488; AP - 1.22, ML 1, DV -4-3.5 mm), amygdala (CTB594; AP -1.3, ML 3.1, DV -4.3), and basal forebrain (CTB647; AP 0.14, ML 1.5, DV -5.5) or septum (CTB647; AP 1.1, ML 0, DV -3-4). One week later, mice were transcardially perfused with PBS and 4% PFA. The brain was removed and left to post-fix in 4% PFA overnight. Next, 50 µm sections were cut, placed onto slides and imaged. Only experiments in which the injection site was restricted to the intended forebrain region were analyzed. Images of the DCN and PBN were imaged on the Zeiss Imager 2 Fluorescent Microscope.

### Retrobead Injections

100 nl of green retrobeads (Lumafluor) were injected into the parabrachial nucleus. Injections were made at a 22 degree angle (parallel to the longitudinal axis of the animal, angled with respect to the dorsal-ventral axis), at coordinates AP -3.725, ML 1.35, and -3.65 mm from the surface of the brain. (**Fig. 2a-c**). This angle of approach allowed us to target the PBN without passing through the cerebellar cortex or the DCN. Retrobeads are an ideal dye to use because their diffusion is quite limited and are thus well-suited to target small brain regions. The use of retrobeads complements previous studies based on viral approaches ^30^. After 3 days, mice were heavily anesthetized, perfused with PBS + 4% PFA, and their cerebellums were removed for histology. A series of sagittal slices (50 μm) were then cut to determine the injection site and PC labelling. Injection sites were scrutinized for leakage into the neighboring vestibular nuclei, or into the cerebellum itself, and mistargeted injections were not included in further analysis. PCs were manually counted in the two slices with the most PC labelling, which were typically 500-600 μm from the midline. Slices were imaged on the Zeiss Imager 2 Fluorescent Microscope.

### Quantification of PC synaptophysin-tdTomato puncta in the brainstem

PCP2-cre mice were crossed with a synaptophysin-tdTomato reporter line to visualize PC boutons. (**Fig. 2d-f, i, Extended Data Fig. 2**). We have used these mice previously to quantify PC synapses, and we found that in addition to the bright tdTomato labelling of boutons, there is less intense labelling of the somata and dendrites, and very faint labelling of the axon ^37,38,70^. Mice were heavily anesthetized, perfused with PBS+4% PFA, and had their brains removed. After post-fixing for one day in PFA, brains were sliced coronally at 50 μm for confocal imaging (Olympus FV1200). Large areas around the PBN (∼2 mm x 2 mm) were imaged under a 60X objective by acquiring multiple overlapping fields and stitching them together in ImageJ’s Grid/Collection stitching plugin. These areas were manually matched and aligned to corresponding DIC brightfield images of the same regions using the cerebellar lobules and the 4^th^ ventricle as guides. The brachium conjunctivum was segmented manually from the DIC image (see **Extended Data Fig. 2**), as was the cerebellum. Puncta from the confocal image were segmented using the MatBots toolbox^71^ in particular the Nuclei Segmentation Bot. In brief, two slices were manually annotated for puncta and used to train a general model to annotate the entire dataset. Once puncta were identified, locations were remapped onto the annotated brightfield image. The model was not trained to differentiate tdTomato in the axon bouton from signals in PC somata and dendrites. Therefore, the cerebellum was not included in further analysis. The brachium conjunctivum was used as a reference point to align and subsequently generate average maps at every anterior-posterior position. All slices were rotated such that the brachium was perfectly horizontal, and the slices were then aligned to the brachium centroids. To use both hemispheres of the brain, maps were flipped such that medial was on the left side of the brachium. Puncta were then binned in 30 μm squares and averaged across corresponding sections. All analysis was conducted in MATLAB.

### Quantification of vGAT and PC axons in the brainstem

To visualize and quantify all inhibitory synapses and inhibitory synapses made by PCs in the brainstem (**Fig. 2gh, Extended Data Fig. 2**), we used an approach similar to that used previously^70^: vGAT immunostaining was used to identify inhibitory synapses and PC axons and boutons were labelled with TdTomato in PCP2-cre;Ai9 mice. We also stained for tyrosine hydroxylase (TH) in these experiments to identify neurons of the LC. Previous studies have reported the presence of PC synapses within the LC ^39^, which is next to the PBN, and these synapses are a potential confound in our studies. Our experiments allowed us to compare the number and density of PC synapses in the LC and the PBN. Mice were anesthetized, perfused with PBS+4% PFA, and had their brains removed. After post-fixing for one day in PFA, brains were sliced coronally (50 μm thick secions). Sections were stained for vGAT (Synaptic Systems, #131 004, 1:500) and tyrosine hydroxylase to identify LC neurons (Immunostar, 22941, 1:1000), and the upper pons was imaged with a confocal microscope (Olympus FV1200) by acquiring 5 μm deep (10 z-steps) stacks of multiple overlapping fields and stitching them together in Imaris Stitcher (Oxford Instruments). Stitched images were imported into Imaris (Oxford Instruments). vGAT puncta were detected using the spot detection feature (1 μm expected spot size). tdTomato labelled axons were detected with the surfaces feature with a fluorescence threshold of 1 standard deviation from the median of the intensity histogram. Overlap of tdTomato and vGAT puncta was defined as zero distance between puncta and surface detection.

To examine the percentage of vGAT overlapping PCP2-cre x tdTomato labelled axons in the LC and the PBN, the borders of each of these structures were manually drawn in 3 dimensions to create 2 3d surfaces using TH labelling and the brachium conjunctivum as guides, respectively. Puncta within each of these surfaces were then quantified. Raw images (**Extended Data Fig. 2**), and analyzed puncta in **Fig. 2gh**, establish that while there are many inhibitory synapses within the LC, a very low number and fraction of these synapses are from PCs. In the PBN, there are many more PC synapses and a much higher fraction of the inhibitory synapses are from PCs.

### Slice Electrophysiology

Slice experiments were performed to provide a functional test of whether PCs directly inhibit PBN neurons (**Fig. 2lm**). Slices were made from adult (>P40) B6.Cg-Tg(Pcp2-cre)3555Jdhu/J x B6;129S-Gt(ROSA)26Sor^tm32(CAG-COP4*H134R/EYFP)Hze^/J (PCP2-cre x ChR2, Jackson Labs) mice. Animals were anesthetized with isoflurane and transcardially perfused with solution composed of in mM: 110 Choline Cl, 2.5 KCl, 1.25 NaH_2_PO_4_, 25 NaHCO_3_, 25 glucose, 0.5 CaCl_2_, 7 MgCl_2_, 3.1 Na Pyruvate, 11.6 Na Ascorbate, 0.002 (R,S)-CPP, 0.005 NBQX, oxygenated with 95% O2/5% CO_2_, and kept at 35°C. 200 µm thick coronal slices were cut (Leica 1200S vibratome) and transferred to a chamber with ACSF (in mM: 127 NaCl, 2.5 KCl, 1.25 NaH_2_PO_4_, 25 NaHCO_3_, 25 glucose, 1.5 CaCl_2_, 1 MgCl_2_). Slices were allowed to recover at 35°C for at least 20 min before experiments.

Borosilicate electrodes (2-4 MΩ) were filled with internal solution (in mM: 110 CsCl, 10 HEPES, 10 TEA-Cl, 1 MgCl_2_, 4 CaCl_2_, 5 EGTA, 20 Cs-BAPTA, 2 QX314, 0.2 D600, pH to 7.3). Experiments were performed at 32-35°C in 5 µM NBQX to block AMPARs, and 2.5 µM (R)-CPP to block NMDARs. Cells were held at -60 mV. PC axons expressing ChR2 were stimulated by 473 nm light from an LED (Thorlabs) through a 60x objective for full-field illumination. Single 1 ms pulses (∼80 mW/mm^2^ steady-state power density as measured under objective) were used. We used a high chloride internal solution to provide good voltage control and highly sensitive recordings^72^. With this internal solution, currents reverse at 0 mV and are inward at a holding potential of -60 mV. The short latency between the laser pulse and onset of the synaptic current (2.3 ms), and the high percentage of PBN neurons that were inhibited by PCs (75 %), established that PCs directly inhibit a large fraction of PBN neurons.

### Implantation of Bilateral Optical Fibers and EMG wires

Following the general surgical protocol (above), two 000-120 × 1/16” screws (Antrin Miniature Specialties, Fallbrook, CA) secured to the skull to stabilize further implants. Two teflon coated tungsten wires (de-insulated 1 mm from implanted end, 100 µm diameter, A-M Systems, Sequim, WA) were implanted superficially under the skin along each side of the mouse’s back for EMG measurements. Then, a custom-made (4.5mm long with 1.7 mm separation) two ferrule dual fiber-optic cannula (Doric lenses, Canada) was implanted at a 22° angle away from the skull at AP -3.72, ML ±1.35, and DV -3.6 for PBN implants or AP -4.72, ML -1.35, and DV -2.9 for DCN implants. Metabond Quick Adhesive Cement (Parkell, Edgewood, NY) was applied to secure the fiber optic and the custom-made titanium head bracket. The mice were then removed from the stereotaxic surgery apparatus, placed in new cage, and allowed to recover for at least 2 days.

### Quantification of c-Fos labelling in the parabrachial nucleus, cerebellum, and forebrain

cFos was used as an activity marker to assess the regions that were activated by the suppression of either the PC-PBN pathway or the PC-DCN pathway (**Fig. 3c-f**) and to determine the downstream regions that were influenced by these pathways (**Fig. 4e-j**). PCP2cre/Halo and wildtype mice were implanted with fiber optics in the parabrachial nucleus or deep cerebellar nuclei as discussed above. Awake head-fixed mice were stimulated with 100 ms pulses unilaterally every 8 seconds for 3 hours. They were then rapidly anesthetized, perfused with PBS+4% PFA, and had their brains removed. After post-fixing for one day in PFA, brains were sliced either coronally or sagittally at 50 μm for imaging. For brains sliced coronally, a DiI coated pin was inserted to indicate the stimulated side. The slices were stained for c-Fos (1:500 dilution, #2250, Cell Signaling) and visualized with the Alexa 647 secondary antibody (1:500 dilution, ab150083, Abcam). Slices of the entire brain were imaged with the whole slide scanner (Olympus VS120). These slide scanner images were then manually matched and aligned to identify the regions noted in **Figs. 3 and 4**. Images were postprocessed in ImageJ using “rolling ball” background subtraction. For display, images were greyscaled and inverted such that the background was white and signal was black. For quantification, two randomly selected areas of 500 μm × 500 μm were chosen in each brain region of interest in six slices from each animal on each side of the brain. Cells were counted blind to condition, and regions were selected using the DAPI channel. The cFos data is summarized in **Table 1, Fig. 3ef Fig. 4lm, Extended Data Fig. 5, and Extended Data Fig. 11**.

### Pupil Dilation, EMG Measurement, and Wheel Movement

Pupils were imaged using a USB camera (Mako U-029B, Edmund Optics) acquiring at 300-330 Hz. The camera was controlled and the pupil diameter was measured online using custom scripts in MATLAB. Signals from implanted wires were amplified using a modified Backyard Brains Spikerbox^73^. Wheel movement was tracked using a disassembled optical mouse and the Arduino Uno microcontroller. Signals were routed to the analog input channels on a RHD2000 recording system (Intan Technologies). EMG signals were then bandpass filtered offline (1-500 Hz) for analysis. Heart beats were isolated first by bandpass filtering the EMG signal (5-200 Hz) and then manually sorting cardiac potentials in Offline Sorter (Plexon).

### Place Preference

Implanted mice were handled by experimenters for at least 15 minutes prior to any procedures to acclimate them to handling. Mice were subsequently acclimated to a Branching Fiber-optic Splitter wire (Doric lenses, Quebec City, QC, Canada) attached to their implanted optical fibers for 1 hour in a 46” by 46” box. The wire was attached to a 1×1 Fiber-optic Rotary Joint (Doric lenses, Quebec City, QC, Canada) which was connected to the 635 nm laser.

Place preference testing occurred in a two chambered box with visual landmarks in each chamber and a doorway between the chambers. This testing chamber was housed in a behavioral testing closet with white noise (60 dB) and illumination at 30 lumens. Place preference testing consisted of 15-minute trials (baseline: no stimulus assigned, test: bottom chamber assigned stimulus, retention: no stimulus assigned, and reversal: top chamber assigned stimulus) that took place over 4 consecutive days. Mice were tracked online with a custom Matlab script and a USB camera (ELP USB with Camera 2.1mm Lens, ELP, China). Continuous 50 ms 10 Hz pulses were delivered when mice entered the assigned stimulus chamber, and ceased when they left. Mice that were found to have a significant initial preference for either chamber in the baseline test (Percent Time > 70%) were excluded from analysis.

### Open Field Experiments

Open field experiments (**Fig. 3m, Extended Data Fig. 7**) were conducted in a subset of mice undergoing the place preference test above. Acclimation and stimulation occurred as above. Tests were done in a behavioral closet with the same white noise (60 dB) and illumination at 30 lumens, and conducted in a square arena in 20 minute trials. Stimulus durations were alternated 1 minute on and 1 minute off. Stimulus trains and intensities were identical to those in the place preference experiments (50 ms 10 Hz pulses). Mice were videotaped with a USB camera (ELP USB with Camera 2.1mm Lens, ELP, China), and tracking was performed offline in a custom Matlab script.

### Confirmation of Implant Locations

PCP2-cre/Halo mice and wildtype were perfused with PBS and 4% PFA. After perfusion, the brains were removed and allowed to post-fix in 4% PFA overnight. After the post-fixing period, the 4% PFA was replaced with PBS. Each brain was sliced sagittally at 100 um in PBS and cover-slipped. All mice with correct bilateral implants in either PBN or DCN were included in the initial analysis shown in **Fig. 3**. We subsequently analyzed mice where only one implant was correctly located in either the PBN or DCN and the other implant was located outside either region, and show that data in **Extended Data Fig. 8**. All bilateral parabrachial nucleus implants and bilateral DCN implants were within the same medial lateral plane (± 100 µm).

### Behavioral Blinding

For the first 36 mice, behavioral testing was carried out blind to genotype (PCP2-cre/Halo or wildtype) and optical fiber implant locations. Wildtypes were not included in the remaining mice, though given that precise optical fiber implant locations were unknown until histology, experiments were blind to condition.

### Quantification and visualization of PC-PBN projections in the forebrain

In order to characterize the projections of PBN neurons that are inhibited by PCs (**Fig. 4a-d, Extended Data Fig. 9**), wildtype mice were injected with 100-200 nl AAV1-Syn-Cre (AAV.hSyn.Cre.WPRE.hGH, Addgene) in the posterior cerebellum (AP -7.2, DV -2-3, ML 0), and 100-200 nl cre-dependent tdTomato (AAV2/1.FLEX.tdTomato.WPRE.SV40, Addgene), and some cases, synaptophysin-YFP (AAV8.2-hEF1a-synaptophysin-EYFP, MGH Vector Core) in the PBN (22 degree angle, AP -7.2, DV-3.65, ML, 1.35). This approach relies on restricting injections of AAV2/1.FLEX.tdTomato.WPRE.SV40 to the PBN. We therefore only characterized mice with injections that were localized to the PBN (**Extended Data Fig. 9**). After waiting two to three weeks, mice were heavily anesthetized, perfused with PBS and 4% PFA, and had their brains removed. After post-fixing for one day in PFA, brains were sliced coronally at 50 μm.

Slices of the entire brain were imaged (Olympus VS120). Images were postprocessed in ImageJ using “rolling ball” background subtraction. Images were manually aligned to the mouse brain atlas using the DAPI channel.

Reconstruction of PBN axons were done for animals also injected with synaptophysin-YFP to confirm the presence of axon terminals in forebrain regions. Fields from the septum, basal forebrain, and amygdala were imaged using a confocal microscope (Olympus FV1200) under a 60X objective with a z-increment of 1 μm.

The Volume Annotation Bot in MatBots^71^ was used to more carefully examine the overlap of synaptophysin-YFP on tdTomato axon signal for several select regions. Multiple slices from the complete 20 μm stack image were manually annotated to train models to automate the reconstructions of the axons across the entire stack. Similar manual annotations were done to identify synaptophysin puncta. Once axons and synaptophysin were segmented, all boutons that did not overlap axons were removed from the 3D reconstructions to more easily visualize axons from cerebellum-recipient parabrachial neurons (**Extended Data Fig. 9**).

### Trans-synaptic labelling of PBN neuronal nuclei

Nuclei in the upper pons putatively receiving posterior cerebellum projections were trans-synaptically labelled for RNA sequencing in the Sun1-GFP mouse. Mice were injected with ∼2000 nl of a 1:6 ratio of AAV1-Syn-Cre (AAV.hSyn.Cre.WPRE.hGH, 105553-AAV1, Addgene) and AAV1-CAG-tdTomato (59462-AAV1, Addgene) in six locations along cerebellum lobule 9. The pipette was angled 10 degrees above horizontal and the injections were 0.5mm above the edge of the skull along two tracts, each +/-0.5 mm lateral to the midline. Along the tracts, ∼300-400 nl was injected at each DV location: -1.2, -0.7, and -0.2 mm. This method is useful for identifying PBN neurons that are directly inhibited by PC neurons because PBN neurons do not project to the posterior vermis ^7,50^. For regions that project to the vermis, such as the LC and the VN^39,53^, the interpretation of labelled cells is more complicated. In addition to anterogradely expressing cre, AAV1-Syn-Cre can produce retrograde expression, albeit with low efficiency ^49^. For that reason, cells in the LC and the VN that are labelled in these experiments could have been labelled as a result of a direct PC connection, or because they project to the vermis. The experiments of **Fig. 5e** address this issue by using a relatively small restricted injection in the posterior vermis that labels the PC fibers with tdTomato, and labels nuclei with AAV1-Syn-Cre. In these experiments labelling within the PBN was restricted to regions with prominent PC fiber labelling. This was not the case for regions outside the cerebellum (LC and VN), which had some labelling in the absence of PC fibers (**Fig. 5e**, *middle*). In the experiments where we were looking for overlap between retrograde labelling from the amygdala and anterograde labelling from the vermis (**Fig. 5fg**), we used larger injections to increase the fraction of anterogradely labelled PBN neurons and increase the chance of overlap. In these experiments we observed more labelled cells, even in regions that we had shown have a low density of PC synapses (**Fig. 5g**). This suggested that retrograde labelling of LC and VN neurons that project to the vermis becomes more prominent for large injections in the vermis. This suggests that for our RNAseq studies, clusters of non-PBN labelled cells may contain retrogradely labelled cells and may not be PC targets.

### Generation of parabrachial nucleus single nuclei profiles

Frozen mouse brains were securely mounted by the frontal cortex onto cryostat chucks with OCT embedding compound such that the entire posterior half including the cerebellum and brainstem were left exposed and thermally unperturbed. Dissection of 500 um anterior-posterior (A-P) spans of each of 10 parabrachial nuclei (bilaterally from 5 mice) was performed by hand in the cryostat using an ophthalmic microscalpel (Feather safety Razor #P-715) and 1 mm disposable biopsy punch (Integra Miltex, York, PA) pre-cooled to -20°C and donning 4x surgical loupes. In order to assess dissection accuracy, 10 μm coronal sections were taken at each 500 μm A-P dissection junction and imaged following Nissl staining. Each excised tissue dissectate was placed into a pre-cooled 0.25 ml PCR tube using pre-cooled forceps and stored at -80°C until dissociation the next day.

Nuclei were isolated from mouse brain samples using a previously published gentle, detergent-based dissociation protocol (dx.doi.org/10.17504/protocols.io.bck6iuze) adapted from one generously provided by the McCarroll lab (Harvard Medical School). See protocols.io link for all buffers and solution concentrations. All steps were performed on ice or cold blocks and all tubes, tips, and plates were precooled for >20 minutes prior to starting isolation. Briefly, the pooled frozen tissue dissectates were placed into a single well of a 12-well plate and 2 ml of ExB was added to the well. Mechanical dissociation was performed through trituration using a P1000 pipette, pipetting 1ml of solution slowly up and down with a 1 ml Rainin (tip #30389212) without creating froth/bubbles a total of 20 times. Tissue was let rest in the buffer for 2 minutes and trituration was repeated. A total of 4-5 rounds of trituration and rest were performed (∼10 minutes). The entire volume of the well was then passed twice through a 26-gauge needle into the same well. Following observation of complete tissue dissociation, ∼2 ml of tissue solution was transferred into a precooled 50 ml falcon tube. The falcon tube was filled with Wash Buffer (WB) to make the total volume 30mls. The 30ml of tissue solution was then split across 2 different 50 ml falcon tubes (∼15 ml of solution in each falcon tube). The tubes were then spun in a precooled swinging bucket centrifuge for 10 minutes, at 600 g at 4°C. Following spin, the majority of supernatant was discarded (∼500 μl remaining with pellet). Tissue solutions from 2 falcon tubes were then pooled into a single tube of ∼1000 μl of concentrated nuclear tissue solution. DAPI was then added to solution at manufacturer’s (ThermoFisher Scientific, #62248) recommended concentration (1:1000).

### Fluorescence-activated nuclei sorting for enrichment of SUN1 positive nuclei

Flow sorting parameters for DAPI gating are described in protocols.io link above. For the SUN1 positive selection on a flow sorter, a DAPI vs 488 gating was established by selecting the 25% highest fluorescent 488 nuclei. Briefly, 0.25ml PCR tube was coated with 5% BSA-DB solution. Solution was then removed and 20 μl of FACS Capture Buffer was added as cushion for nuclei during sort. Nuclei were sorted into a chilled 96 well FACS plate (Sony M800 FACSorter) holding the 0.25 ml PCR tube. Sorting was done at a pressure of 6-7, with forward scatter gain of 1% on DAPI gate. The “purity” mode was used, and no centrifugation was performed after flow sorting of nuclei. Following sort, SUN1 positive nuclei (9.2% of total) were counted using a hemocytometer and adjusted to an appropriate concentration before loading 5825 nuclei into a single lane of the 10x Genomics (Pleasanton, CA) Chromium v3.1 single cell 3’ expression analysis system. Reverse transcription, indexed library generation, and sequencing were performed according to the manufacturer’s protocol.

### Analysis of single nuclei profiles

A pre-filtered Digital Expression Matrix (DGE) was imported in RStudio (R version 4.1.2) and data were analyzed using Seurat package (version 3.2.3). Briefly, we filtered out nuclei that: have a mitochondrial percent >10 and a gene count <200. In addition, we retained only genes that were expressed at least in 10 nuclei across the entire dataset. Even though our viral strategy targetted only neuronal nuclei, we could not avoid a small glial contamination due to the inability of the FACS sorting to have an efficiency of separation of 100%. Therefore, the first clustering, containing a small percentage of glial nuclei, allowed us to exclude them from downstream analyses. After filtering out low quality nuclei and glial contamination, we ended up with a dataset of 3,876 neuronal nuclei × 15,479 genes. The following Seurat functions were used: 1. CellCycleScoring() to calculate quantitative scores for G2M and S phase. Briefly, this function assigns each cell a score, based on the expression of G2/M and S phase markers list ^74^, and classifies the cell in one of the 3 phases: G1, G2/M and S; 2. SCTransform() to normalize, scale, identify the top 3,000 variable features, and to regress out for the mt.percent and cell cycle covariates; 3. runPCA() to calculate the first 50 Principal Components (PCs) of the 3,000 most variable features; 4. runTSNE() a dimensional reduction method, to visualize cell clusters; 5. FindNeighbors() to construct a Shared Nearest Neighbor (SNN) Graph, which determines the k-nearest neighbors of each cell and then uses this knn graph to construct the SNN graph by calculating the neighborhood overlap (Jaccard index) between every cell and its k.param nearest neighbors; 6. FindClusters() to define cell clusters and their granularity using the original Louvain algorithm at resolution of 0.8. 7. FindAllMarkers() to perform a differential gene expression between each single cluster and all the others using the non-parametric Wilcoxon Rank Sum statistics. A gene was considered differentially expressed if the average logFC was >0.5 and had a Bonferroni adjusted P-value <0.01.

### Combined retrograde and anterograde labelling of PBN neurons receiving PC input

To visualize PBN neurons projecting to the forebrain and receiving input from PCs, we used the same trans-synaptic labelling strategy as above, except in the Ai75, nuclear-localized tdTomato mouse for better signal to noise. We combined this with cholera toxin (CTB647 as in **Fig. 1**) injected in the amygdala. After two weeks, mice were perfused, fixed, and had their brains removed. They were sliced into 50 μm sections. Regions around the PBN were imaged using a confocal microscope (Leica Stellaris X5, 20X objective with 15-20 μm stacks at 1 μm steps). tdTomato nuclei were detected using custom scripts in Matlab. CTB fluorescence was dense and often dominated by processes, and it was difficult to definitively label CTB labelled cells. Instead, tdTomato nuclei were examined for colabelled CTB, first by automatically thresholding for CTB in the immediate vicinity of each tdT nuclei, and then by manually inspecting each putatively colabelled nuclei to confirm the colabel. Annotated sections were aligned to the center of the brachium conjunctivum. Four such sections (8 total PBN) were annotated and were qualitatively similar. Displayed in **Fig. 5g** is a combination of two such annotated sections.

### Immunohistochemistry and fluorescence in situ hybridization (FISH) following trans-synaptic virus injection

Injected Sun1GFP mice were perfused, fixed, and had their brains removed. They were then sliced into 50 μm sections and stained with anti-FoxP2 (1:500, Abcam, ab16046) and anti-GFP (1:1000, Abcam, ab13970). 8 μm stacks were imaged using a confocal microscope (Olympus, FV1200), and further analysis was done in Matlab (Mathworks). Images were processed and thresholded such that Sun1-GFP positive nuclei were unambiguously registered. Because FoxP2 positive soma were often embedded within FoxP2 expressing processes, we found it sufficiently ambiguous that we could not rely on automated approaches to identify FoxP2 somas. Instead, we manually inspected every Sun1-GFP positive nuclei/ROI for FoxP2 positive somas to generate the resulting map in **Fig. 5**.

For FISH experiments, we switched from the Sun1GFP mouse to the Ai75, nuclear-localized tdTomato mouse for compatibility with FISH probes. All injections in the Ai75 mice were done identically except that the AAV1-CAG-tdTomato virus was omitted. Brains from injected Ai75D mice were rapidly removed, frozen, and embedded in optimal cutting temperature (OCT) compound (Tissue-Tek). Tissue was cut on a cryostat (Microm HM500-CM) at a thickness of 20 μm. Fluorescent in situ hybridization was performed according manufacturers protocols (ACD-Bio RNAscope Multiplex Assay manual, document Number 320513), using fluorophore-conjugated probes, Lmx1a-C1 probe (Cat#493131, ACD-Bio, CA), tdTomato-C2 (Cat#317041, ACD-Bio, CA), and Pax5-C3 (Cat#311481, ACD-Bio, CA). Slices were then imaged using whole slide scanning microscope (Olympus VS120) with a 40X objective at three different planes separated by 2 μm. Images were processed in Matlab (Mathworks). Briefly, stacks were collapsed into a single plane (max projection), and backgrounds were removed by subtracting the median filtered from the original image. A threshold was applied and single puncta (RNAs) were registered. Clusters of signal were identified by determining which puncta had at least 8 neighbors within a radius of 10 μm. Unclustered puncta were not plotted for visualization.

## Data Availability

Analysis relating to the RNA-seq dataset has been deposited on Zenodo (DOI:10.5281/zenodo.6653204). Other data that support the findings of this study are available from the corresponding author upon reasonable request.

## Author Contributions

C.H.C. and W.G.R. conceived of the project. C.H.C, L.N.N., A.P.S., K.E.B, and W.G.R designed experiments. C.H.C., L.N.N., A.P.S. and D.Z. performed and analyzed *in vivo* electrophysiology experiments. C.H.C. performed *in vitro* slice electrophysiology experiments. C.H.C., A.P.S., L.N.N., Z.Y., and I.F. performed and analyzed anatomical/histological experiments. C.H.C., A.P.S., Z.Y. and K.E.B conducted and analyzed c-Fos functional anatomy experiments. C.H.C, K.E.B., K.M., Z.Y. and L.N.N. performed and analyzed behavioral experiments. C.H.C, A.P.S., and K.E.B, prepared samples for RNA-seq experiments. C.V. and N.N. isolated and processed samples for RNA-seq, and S.N. analyzed. C.H.C. and W.G.R wrote the paper with inputs from all authors.

## Acknowledgements

We thank members of the Regehr lab and David Ginty for comments on the manuscript. In particular we thank Brielle Miano for assistance with RNAscope experiments. This work was supported by grants from the NIH, R01NS032405 and R35NS097284 to W.G.R, K99NS110978 and F32NS101889 to C.H.C., R01DK075632 to B.B.L., and NINDS P30 Core Center (NS072030) to the Neurobiology Imaging Center at Harvard Medical School.

## Competing Interests

The authors declare no competing interests

## Notes

### Competing Interest Statement

The authors have declared no competing interest.

### Summary of Updates

New experiments were performed and Figure 2gh, Figure 3efm, Figure 4lm, Figure 5, Extended Data Figures 2, 9, 10 and 12, and Tables 5, 6 and 7 were added. The text has been revised extensively. New authors have been added.

