## Supplemental Figures and Tables for "A Purkinje cell to parabrachial nucleus pathway enables broad cerebellar influence over the forebrain"

**Extended Data Figures and Tables**

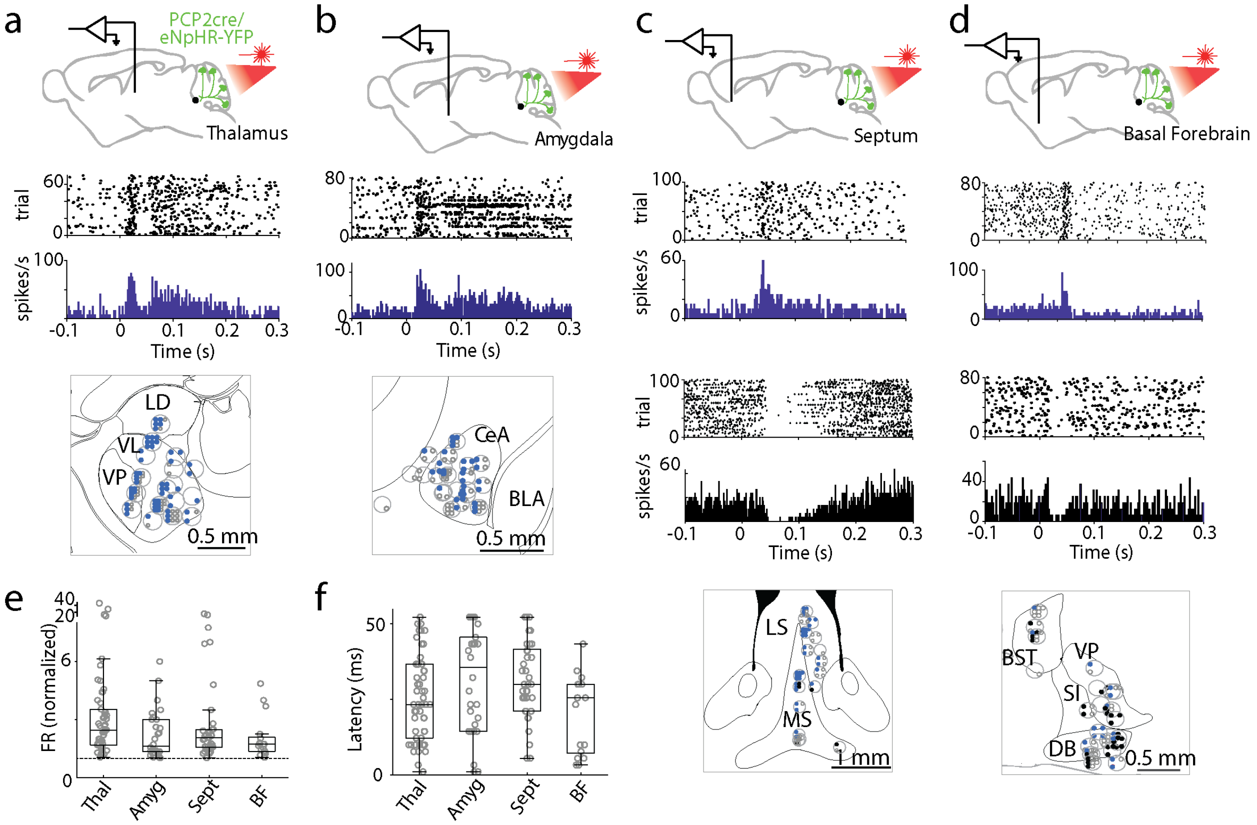


**Extended Data Fig. 1: Suppressing PC firing evoked short latency responses in multiple brain regions.** Single-unit, multielectrode array recordings were made from awake, head-restrained PCP2Cre/Halo mice across several areas in the brain. Example single cell responses are shown for the thalamus (**a**), amygdala (**b**), septum (**c**), and basal forebrain (**d**). Pauses in activity after stimulation were observed in the septum and basal forebrain in a fraction of recorded cells. Example pauses are shown in black. Recording sites are indicated in the grey circles within the corresponding panels on the lower panels. Blue dots indicate responding cells at each site, black dots indicated cells with pauses in firing after stimulation, and open circles indicate nonresponding cells. **e, f.** Box plots representing normalized change in firing rate (**e**) and average latency (**f**) for each region.
LD: laterodorsal, VL: ventrolateral, VP: ventral posterior, CeA: central amygdala, BLA: basolateral amygdala, LS: Lateral septum, MS: medial septum, BST: bed nucleus of the stria terminalis, VP: ventral pallidum, SI: substantia innominate, and DB: diagonal band of broca.

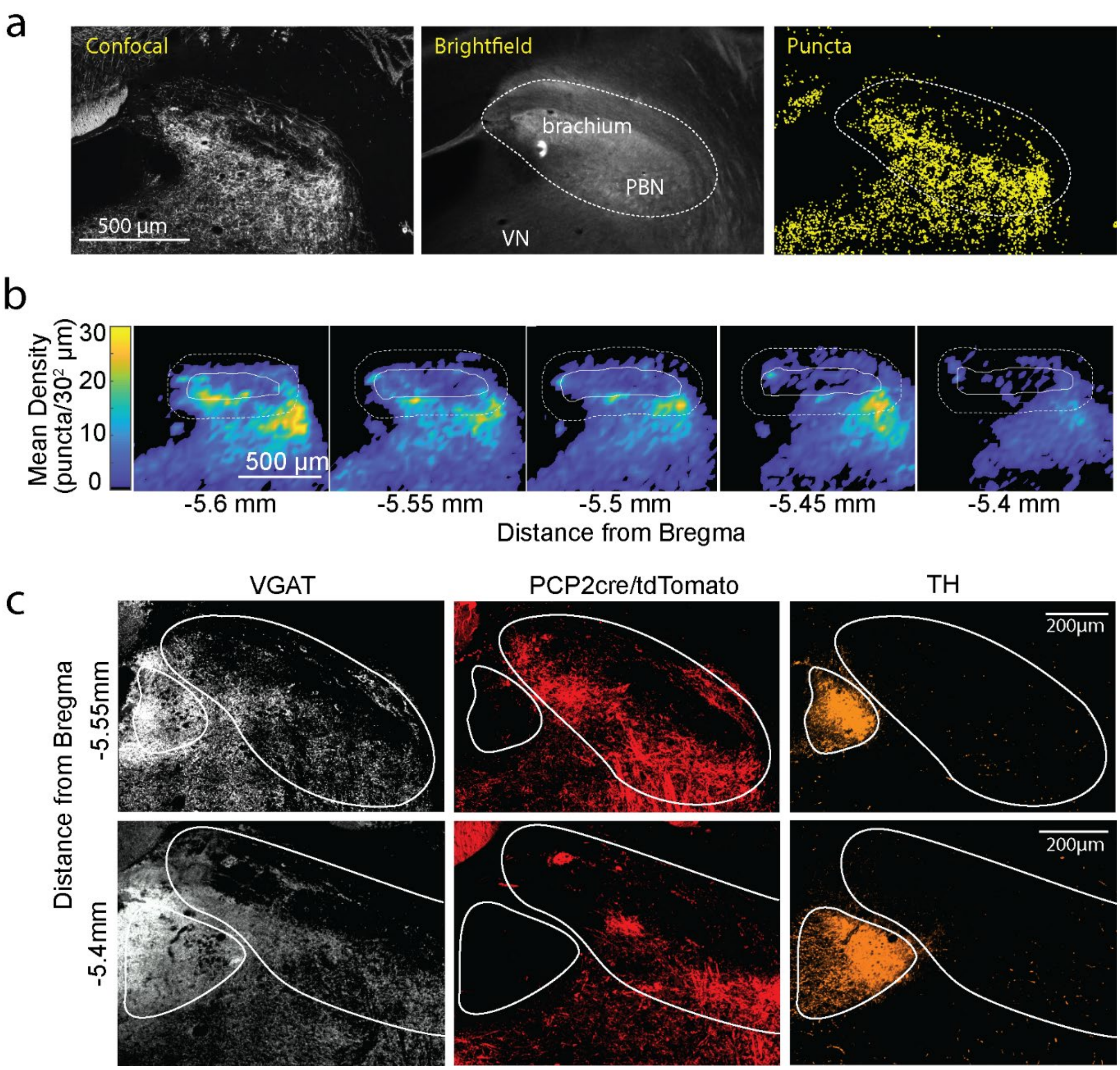


**Extended Data Fig. 2: Purkinje cell synapses within the PBN**

**a.** Method for registering PC synapses and identifying the PBN in the PCP2cre/Synaptophysin-tdtomato mouse. The confocal image (left) showing tdTomato fluorescence, the brightfield image (middle) with the brachium conjunctivum noted, and identified presynaptic boutons (right, yellow) are shown with the bounds of the PBN delineated.

**b.** Average heatmaps of identified PC synaptophysin-tdT puncta in the PBN. Individual slices were binned at $30^2$ µm and averaged by aligning the center of the brachium. Each heatmap is an average of 4 slices (from 2 animals, 2 hemispheres per animal). The bold white line outlines the brachium, and the dotted line the PBN.

**c.** Immunohistochemistry was used to identify inhibitory synapses (vGAT, *white*), and the locus coeruleus (tyrosine hydroxylase, *orange*) in a PCP2cre/tdTomato mouse where PC axons and boutons were labelled (tdT, *red*). Very few PC boutons were apparent in the locus coeruleus.

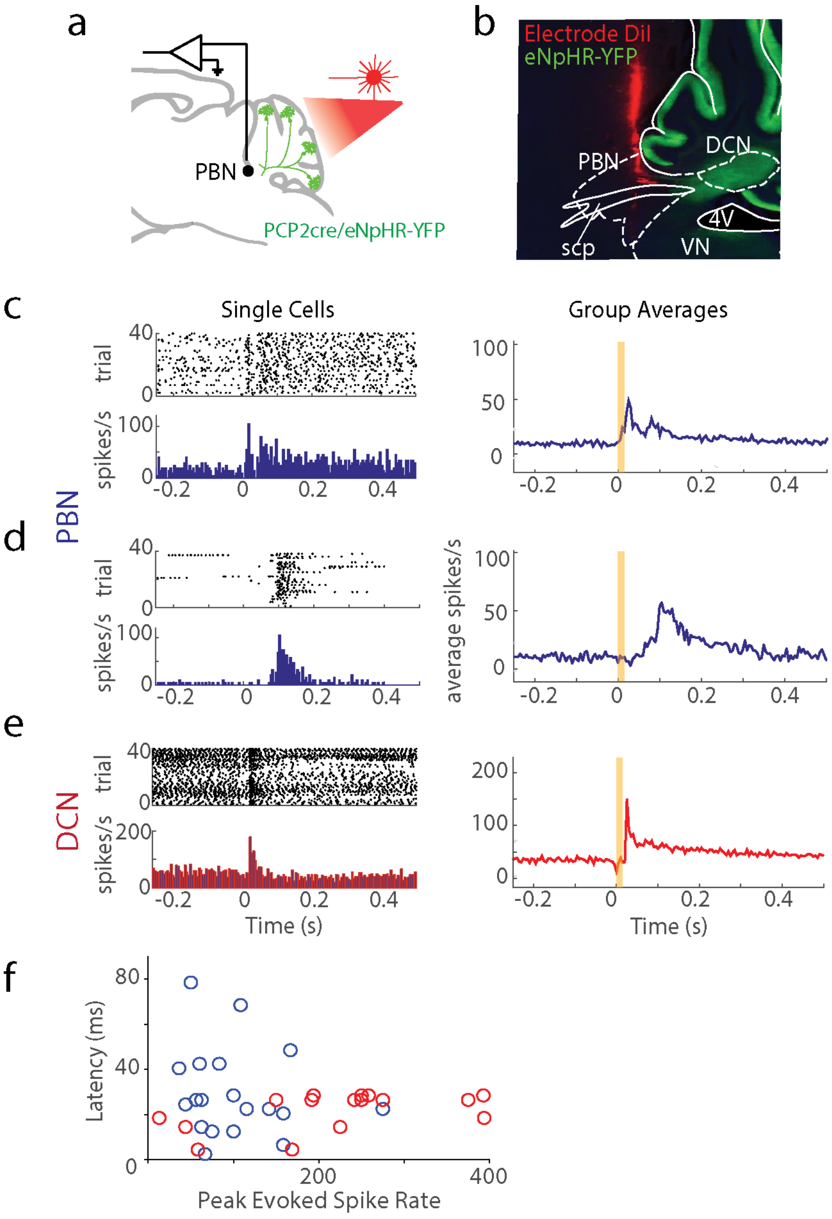


**Extended Data Fig. 3: PBN neurons rapidly increase firing in response to suppression of PC firing.**

- **a.** Single-unit, multielectrode array recordings were made in the PBN or DCN in awake, head-restrained PCP2cre/Halo mice (n=6). The posterior cerebellar cortex was stimulated through a thinned skull (20 ms, red light).
- **b.** Recording sites were recovered by coating the silicon probe with DiI. An example electrode tract is shown. scp: superior cerebellar peduncle, 4V: 4$^{th}$ ventricle
- **c.** *left***,** Firing evoked in a rapidly responding PBN neuron (<30 ms latency).
  *right*, Summary of rapidly responding PBN neurons (13/28 neurons).
- **d.** Same as C but for slower responding PBN neurons (6/28 neurons).
- **e.** Same as C but for DCN neurons (16/16 neurons).
- **f.** Latencies of PBN neurons (blue) and DCN neurons (red) as a function of evoked firing rate.

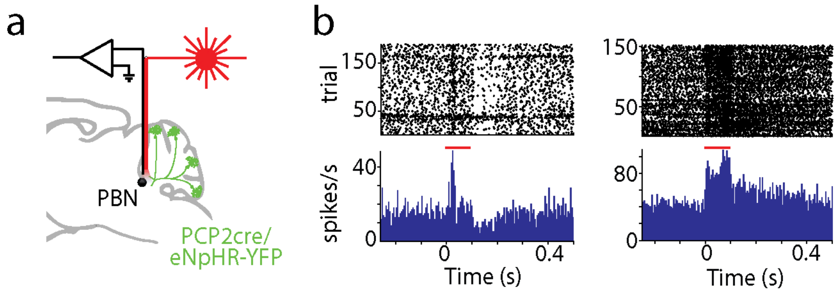


**Extended Data Fig. 4: Suppressing the PC-PBN pathway increases firing in PBN neurons.**

**a.** Awake, head-restrained single-unit recordings in the PBN were made using an optrode in a Halo/PCP-Cre mouse.

**b.** Two example cells showing increases in firing during the 100 ms light pulse (*red line*).

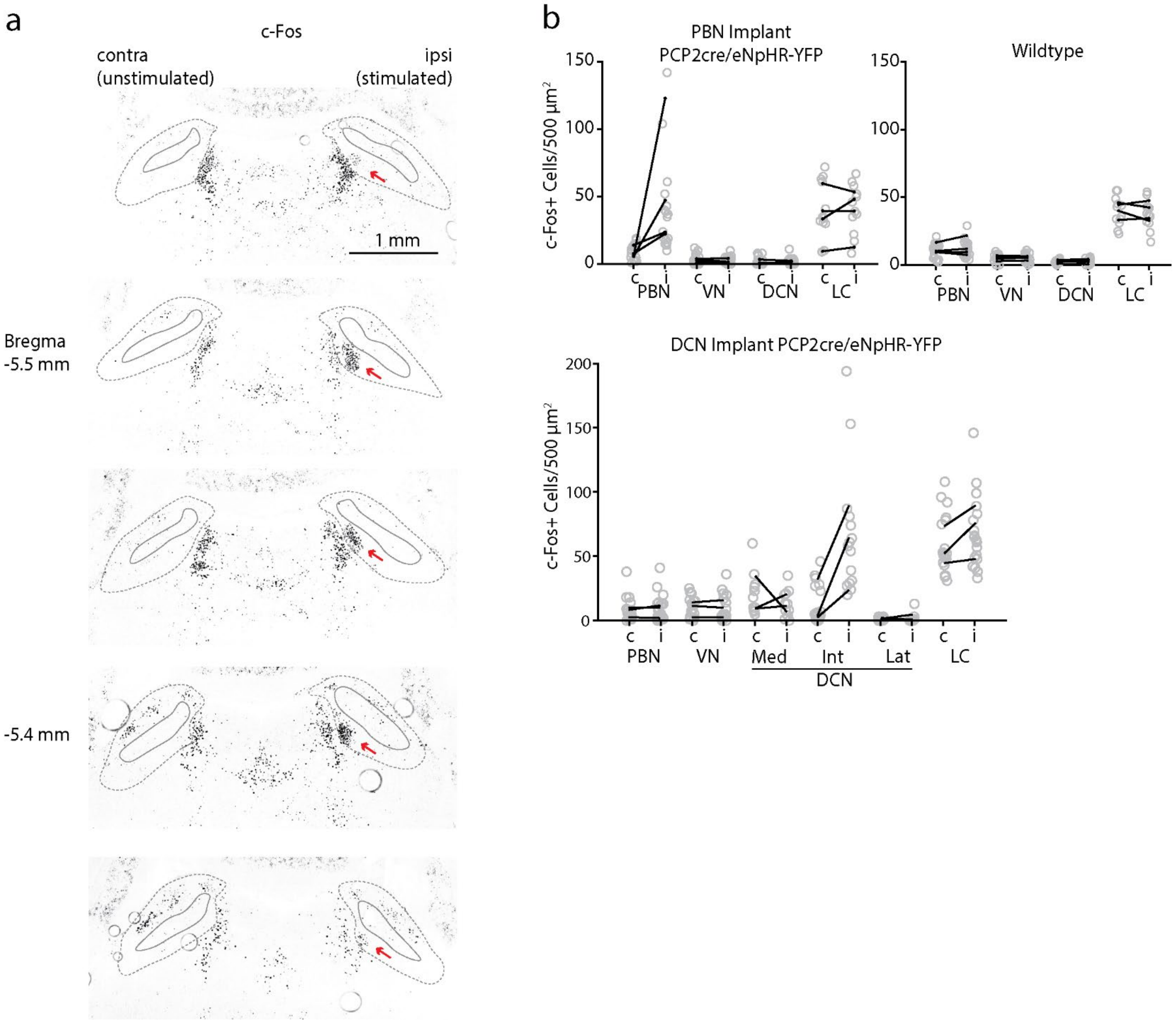


**Extended Data Fig. 5: Specificity of suppression of the PC-PBN pathway.**

- **a.** Images for an example c-Fos stimulation experiment shown in **Fig. 3**. PC inputs to the PBN were suppressed by activating halorhodopsin. In this experiment, unilateral c-Fos upregulation was observed in medial side of the PBN near the tip of the optical fiber(*red arrow*). Brachium is indicated in the solid grey lines, and bounds of the PBN in dotted lines.
- **b.** Hindbrain c-Fos densities for PC-PBN suppression (*upper left*), wildtype (*upper right*), and PC-DCN stimulated (*bottom*) animals. Average numbers of c-Fos+ cells per mouse plotted in black lines for the stimulated (i) and unstimulated (c) sides. Individual areas plotted in grey circles.

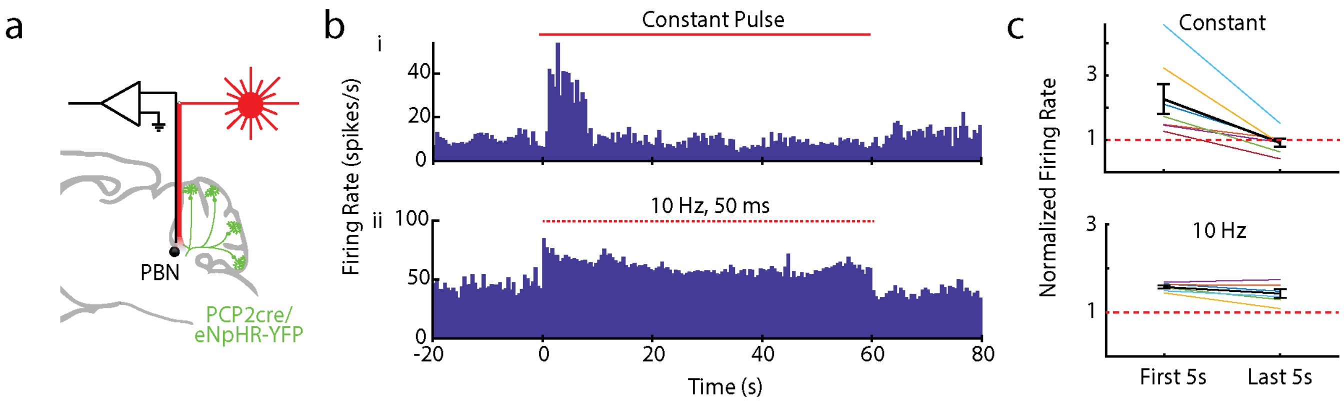


**Extended Data Fig. 6: Suppression of PC-PBN axons can stably increase the firing rates of PBN neurons.**

**a.** Recordings were made from PBN neurons with an optrode. Light was delivered to the immediate region with the attached optical fiber.

**b.** i. Example single cell response to continuous light delivered to PC-PBN axons.
ii. Example response of another cell to pulsed (10 Hz, 50 ms pulses) light delivered to PC-PBN axons

**c.** Summary data showing the firing rate in the first and last 5 s of the train for continuous (top) or pulsed (bottom) trains. We used 10 Hz, 50 ms pulses to suppress the PC-PBN in behavioral experiments due to the superior stability of PBN neuron firing.

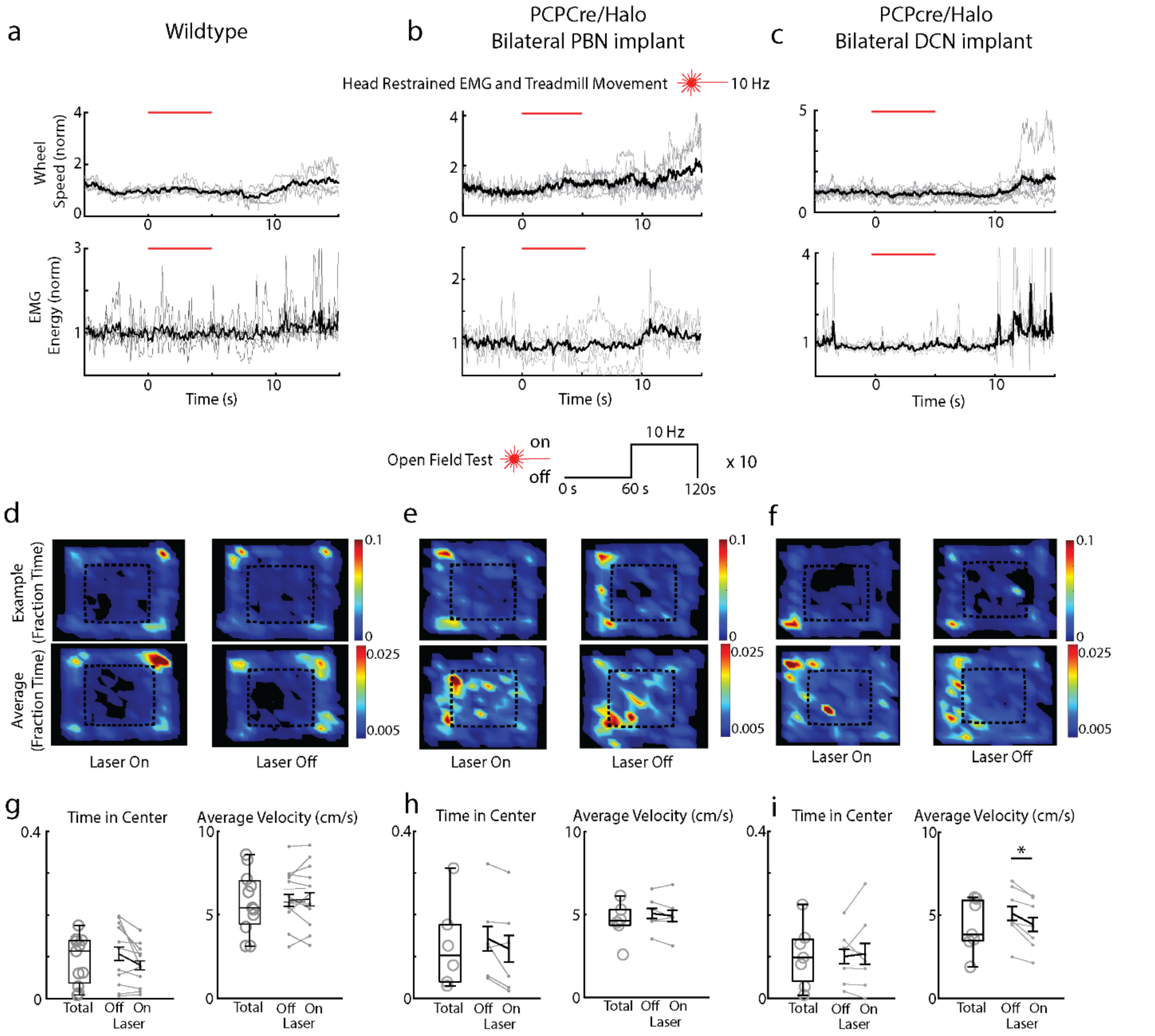


**Extended Data Fig. 7: Effects of PC-PBN and PC-DCN suppression on motor behaviors**

**a-c.** Halo/PCP-cre and wildtype control mice with bilateral optical fiber implants targeting their PBN or DCN were subcutaneously implanted with wires in their backs to measure a field EMG (n = 5). Mice were head restrained over a freely moving wheel. Speed of the wheel (top) and field EMG (bottom) during stimulation (10 Hz, 5 s), were measured (n = 5). (PBN implant EMG/Wheel n=5/5, DCN implant EMG/Wheel n = 4/7, control animals EMG/Wheel n=5/5).

**d-f.** Mice in those groups were also tested in the open field, where 10 Hz optical stimuli were delivered in 60 s intervals. Average position heat maps are shown for each interval (stimulus on and stimulus off).

**g.** Fraction of time in center and average velocity during the open field for the total duration of the test and for the durations while the stimulus was off and on for wildtype control animals

**h.** As in **g**, but for PCP2cre/Halo animals implanted in the PBN.

**i.** As in **h**, but for PCP2cre/Halo animals implanted in the DCN.

* $p<0.05$, see **Table 4**

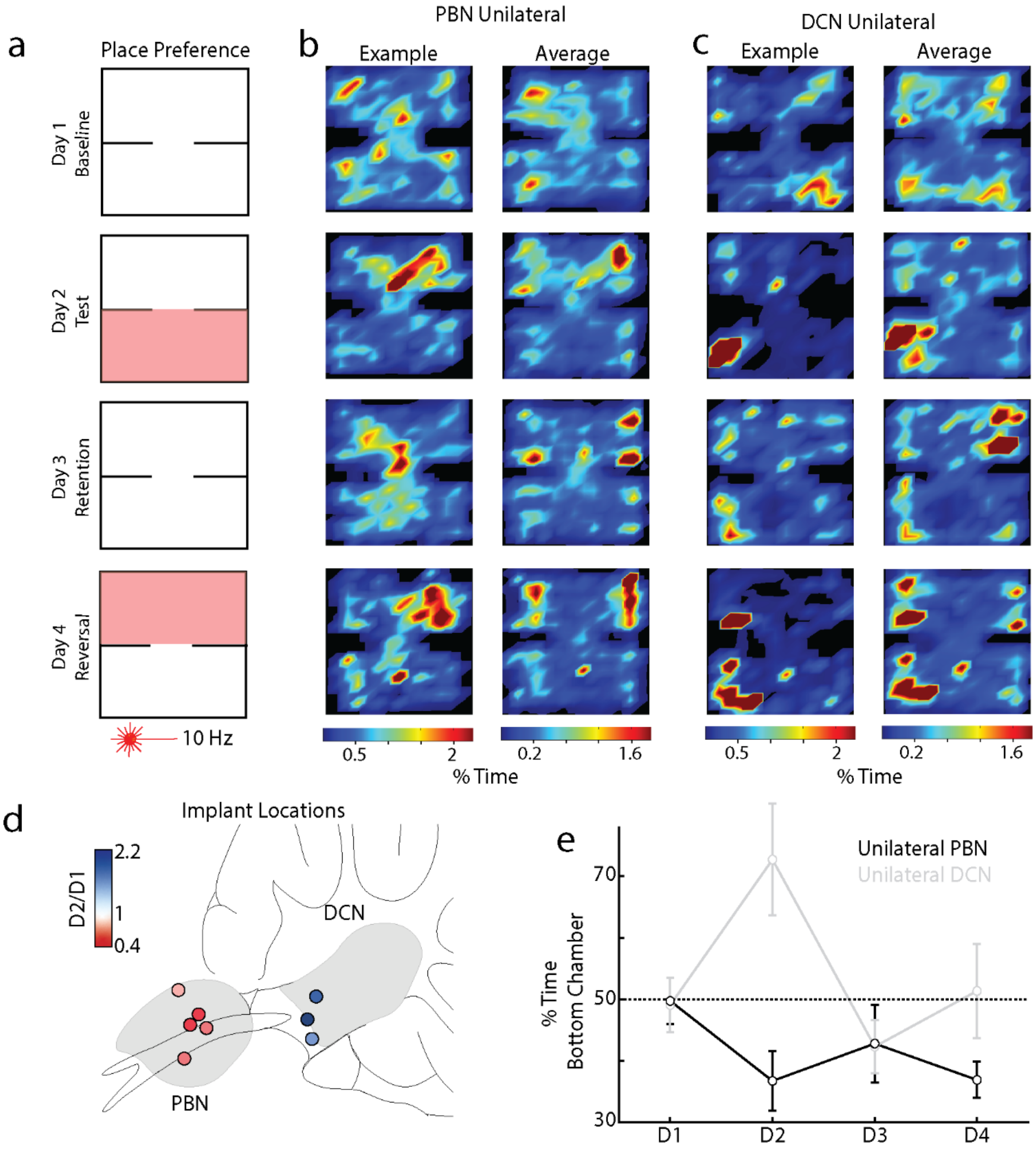


**Extended Data Fig. 8: Place preference for animals with one optical fiber implanted correctly**

- **a.** Attempted bilateral implants often resulted in one optical fiber implanted in a target region, and another implanted completely elsewhere devoid of halorhodopsin expression. We examined these mice's behavior in the place preference protocol as in **Fig. 3**
- **b.** Example (left) and average (right) position heatmaps for corresponding days of the place preference protocol for mice with unilateral implants in the PBN (**B,** n=5).
- **c.** As in **b**, but for unilateral implants in the DCN (**C**, n=3)
- **d.** Implant locations for all mice with color indicating bottom chamber preference (Test/Baseline; D2/D1).
- **e.** Summary of the % time spent in the bottom chamber across all test days for PBN (red) and DCN (blue) unilaterally implanted PCP2cre/Halo mice. (see **Table 3**)

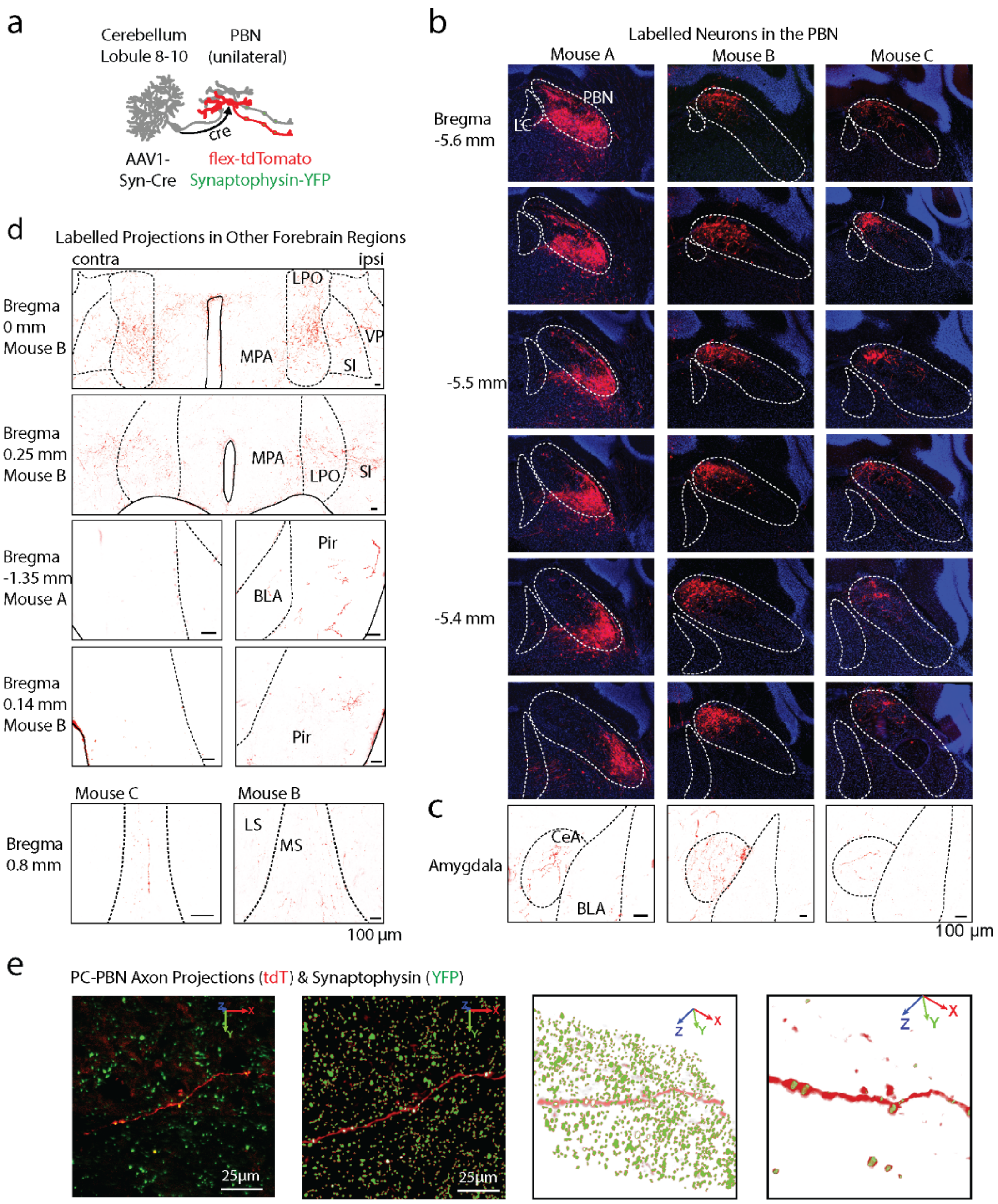


**Extended Data Fig. 9. Labelling of the PC-PBN projection pathway.**

- **a.** An anterograde AAV-cre was injected into the posterior cerebellar vermis, and AAVs with cre-dependent tdTomato were injected into the PBN. This led to tdT-expression in PC-recipient PBN neurons. The PBN of mouse C was also injected with AAV1-Synaptophysin-YFP to label the presynaptic boutons of all PBN neurons.
- **b.** A series of sections shows that tdT expression was restricted to the PBN of the three mice injected as in A.
- **c.** tdT-expressing axons are shown in the amygdala of each mouse.
- **d.** tdT-expressing axons are shown in the indicated regions for the indicated mice.
- **e.** To visualize PC-PBN projections and determine synaptophysin overlap, 20 µm confocal stacks of each region were taken (top panel), and synaptophysin and axon signals were segmented out (middle panels). Puncta associated with a tdTomato-expressing axon are displayed in the examples for the indicated regions.

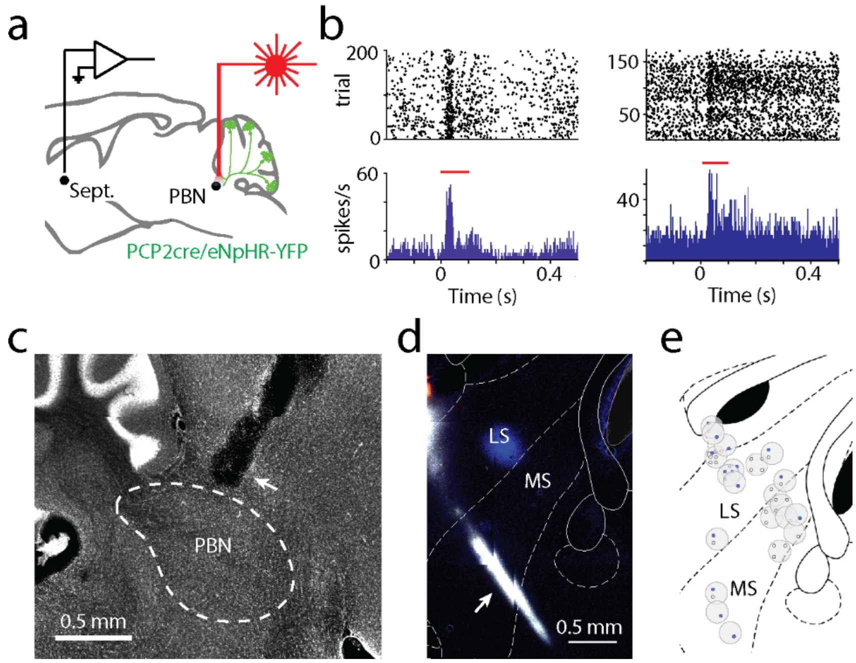


**Extended Data Fig. 10: Suppressing the PC-PBN pathway rapidly increases firing in the septum**

- **a.** In awake, head-restrained mice expressing halorhodopsin in PCs, the PC-PBN pathway was optically suppressed and single-unit recordings were made in the septum.
- **b.** Two example cells showing increases in firing during the 100 ms light pulse (*red line*).
- **c.** Optical fiber implant into the parabrachial nuclei. Shown is a sagittal slice of the PBN with cerebellum on the left. Optical fiber implant is indicated with the white arrow. Boundaries of the PBN are indicated with white dotted lines.
- **d.** Example recording site through the medial septum (MS) and lateral septum (LS). Probes were coated with different color dyes to enable post-hoc verification of recording sites. Recording sites were determined by matching the dye with the final position of the probe inside the brain. Dye is shown in white.
- **e.** Recording sites for all experiments shown above. Every grey circle indicates a recording site. Blue dots indicate responding cells and black dots indicate nonresponding cells for each site.

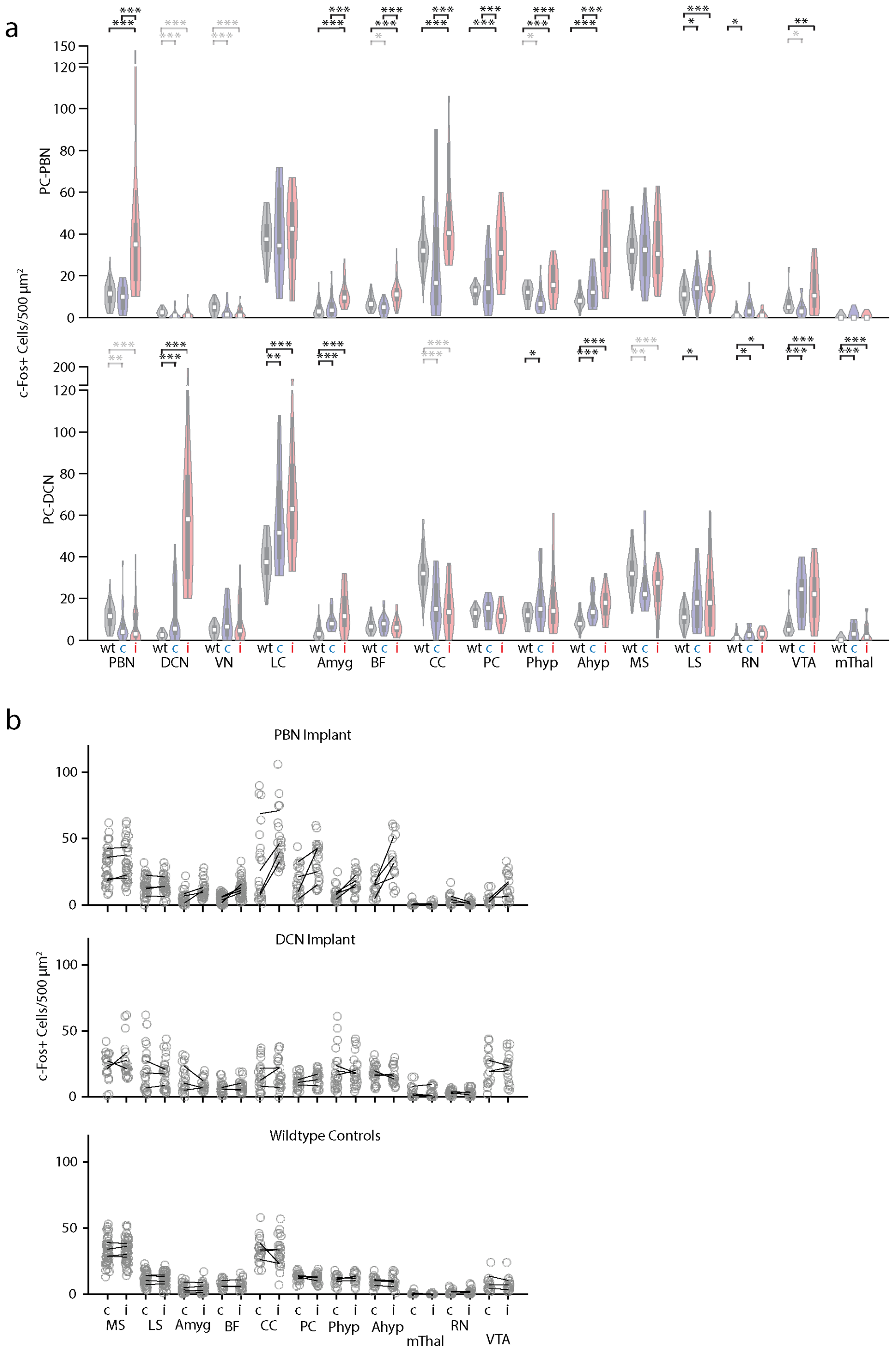


**Extended Data Fig. 11: Upregulation of c-Fos following unilateral suppression of the PC-PBN pathway.**

**a.** Distributions of c-Fos positive cells following inhibition of PC-PBN or PC-DCN pathways. Data corresponds to that shown in **Fig. 3a-f** and **Fig. 4h-m**. For wildtype mice hemispheres were combined because there was no apparent difference between hemispheres (see **B**). * $p<0.05$, ** $p<0.01$, *** $p<0.001$. Grey indicates a decrease in the highlighted comparison. “c” and “I” correspond to contralateral and ipsilateral.

**b.** Corresponding regions and c-Fos densities for PC-PBN stimulated (top), PC-DCN stimulated (middle), and wildtype (bottom) animals. The average number of c-Fos+ cells in different regions are plotted for each mouse (*black lines*). Cell counts are plotted for individual slices (*grey symbols*). See **Table 1.**

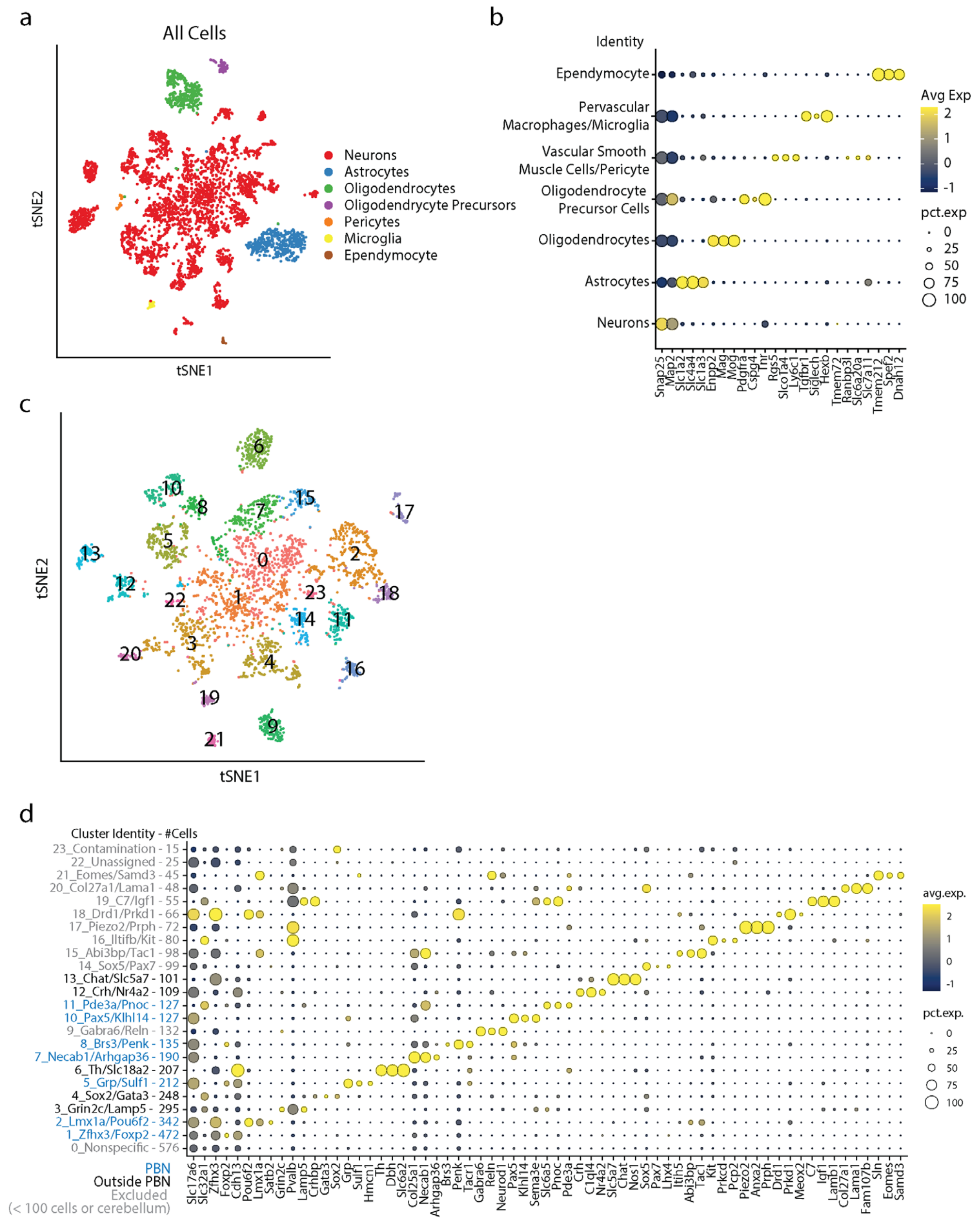


**Extended Data Fig. 12: RNAseq of brainstem neurons labelled following injections of AAV1-Syn-Cre into the posterior cerebellum of CAG-Sun1/sfGFP mice (Fig. 5a).**

- **a.** tSNE visualization of all cells following fluorescence activated nuclei sorting. Clusters were sorted into cell types based off of expression of markers indicated in **b**.
- **b.** Dot plot of scaled expression for indicated clusters
- **c.** tSNE visualization of just neurons from **a.** Cluster numbers correspond with those in **d**.
- **d.** Dot plot of scaled expression for indicated clusters ordered from largest to smallest, with number of cells in each cluster indicated in the axis label. Shown are 3 of the most significantly expressed genes for each cluster, along with Slc17a6 (glutamatergic neurons) and Slc32a1 (GABAergic neurons). Clusters with < 100 cells were categorically excluded. Cluster 9 contains Gabra6, a known marker for cerebellar granule cells, thus likely reflects cerebellar contamination, and was excluded from subsequent analysis.

**Table 1: c-Fos Statistics**

| Figure | ROIs | Animals | Average (mean) | S.E.M. | Statistics (w=ranks) |
|---|---|---|---|---|---|
| **Fig 3E+F & Extended Data Fig 5B:** | | | | | |
| **PBN (halo PBN implant)** | Contra: 16<br>Ipsi: 17 | 4 | 9.5<br>40.5 | 1.5<br>8.5 | One-sided Wilcoxon rank sum<br>p=1.44e-5; w=155.5 |
| **PBN (halo DCN implant)** | Contra: 24<br>Ipsi: 24 | 3 | 7.0<br>8.0 | 1.7<br>2.0 | Wilcoxon rank sum<br>p=0.6334; w=611.5 |
| **PBN (wt)** | 42 total (21/21 contra/ipsi) | 4 | 11 | 0.9 | Wilcoxon rank sum<br>vs. contra PBN p=0.3928; w=1289<br>vs. ipsi PBN p=8.2e-7; w=965.5<br>vs. contra DCN p=6.83e-4; w=549.5<br>vs. ipsi DCN p=0.005; w=594 |
| **VN (halo PBN implant)** | Contra: 26<br>Ipsi: 45 | 4 | 2.7<br>2.2 | 0.6<br>0.3 | One-sided Wilcoxon rank sum<br>p = 0.5797; w = 952 |
| **VN (halo DCN implant)** | Contra: 18<br>Ipsi: 18 | 3 | 9.3<br>9.4 | 2.0<br>2.4 | One-sided Wilcoxon rank sum<br>p=0.75; w=322.5 |
| **VN (wt)** | 32 total (16/16 contra/ipsi) | 4 | 5.0 | 0.5 | Wilcoxon rank sum<br>vs. contra PBN p=0.003; w=1135<br>vs. ipsi PBN p=2.8e-5; w=1650<br>vs. contra DCN p=0.17; w=527<br>vs. ipsi DCN p=0.62; w=483.5 |
| **DCN (halo PBN implant)** | Contra: 30<br>Ipsi: 40 | 4 | 1.6<br>1.5 | 0.5<br>0.3 | One-sided Wilcoxon rank sum<br>p = 0.1641; w = 986 |
| **DCN-Interposed (halo DCN implant)** | Contra: 16<br>Ipsi: 15 | 3 | 14.1<br>66.4 | 3.8<br>12.7 | One-sided Wilcoxon rank sum<br>p = 6.3e-5; w = 158.5 |
| **DCN (wt)** | 32 total (16/16 contra/ipsi) | 4 | 2.5 | 0.3 | Wilcoxon rank sum<br>vs. contra PBN p=0.003; w=1214.5<br>vs. ipsi PBN p=1.5e-3; w=1440.5<br>vs. contra DCN p=0.006; w=517.5<br>vs. ipsi DCN p=3.9e-8; w=600 |
| **LC (halo PBN implant)** | Contra: 12<br>Ipsi: 12 | 4 | 40.2<br>41.0 | 6.0<br>5.3 | One-sided Wilcoxon rank sum<br>p=0.42; w=146 |

| | | | | | |
|---|---|---|---|---|---|
| **LC (halo DCN implant)** | Contra: 20<br>Ipsi: 21 | 3 | 57.4<br>71.1 | 5.1<br>6.9 | One-sided Wilcoxon rank sum<br>p=0.07; w=498 |
| **LC (wt)** | 24 total (12/12 contra/ipsi) | 4 | 38.5 | 2.1 | Wilcoxon rank sum<br>vs. contra PBN p=0.46; w=1214.5<br>vs. ipsi PBN p=0.95; w=446.5<br>vs. contra DCN p=0.004; w=571.5<br>vs. ipsi DCN p=1.88e-5; w=671.5 |
| **Fig 4L+M & Extended Data Fig 11:** | | | | | |
| **Medial Septum (halo PBN implant)** | Contra: 36<br>Ipsi: 34 | 4 | 30.6<br>33.3 | 2.3<br>2.5 | One-sided Wilcoxon rank sum<br>p = 0.26; w=1264 |
| **Medial Septum (halo DCN implant)** | Contra: 20<br>Ipsi: 20 | 3 | 27.6<br>24.2 | 3.3<br>2.4 | Wilcoxon rank sum<br>p=0.96; w=407.5 |
| **Medial Septum (wildtype)** | 96 total (48/48 contra/ipsi) | 4 | 32.9 | 0.9 | Wilcoxon rank sum<br>vs. contra PBN p=0.29; w=2188<br>vs. ipsi PBN p=0.77; w=2171<br>vs. contra DCN p=0.0026; w=758.5<br>vs. ipsi DCN p=0.0025; w=756 |
| **Lateral Septum (halo PBN implant)** | Contra: 40<br>Ipsi: 42 | 4 | 14.3<br>15.3 | 1.1<br>1.0 | One-sided Wilcoxon rank sum<br>p=0.29; w=1804 |
| **Lateral Septum (halo DCN implant)** | Contra: 26<br>Ipsi: 25 | 3 | 17.4<br>20.3 | 2.4<br>3.3 | Wilcoxon rank sum<br>P=0.81; w=663.5 |
| **Lateral Septum (wildtype)** | 96 total (48/48 contra/ipsi) | 4 | 11.5 | 0.5 | Wilcoxon rank sum<br>vs. contra PBN p=0.02; w=3168<br>vs. ipsi PBN p=6.8e-4; w=3597<br>vs. contra DCN p=0.048; w=1883.5<br>vs. ipsi DCN p=0.06; w=1783.5 |
| **Amygdala (halo PBN implant)** | Contra: 28<br>Ipsi: 32 | 3 | 5.0<br>10.9 | 1.0<br>1.1 | One-sided Wilcoxon rank sum<br>p=7.53e-6; w=562 |
| **Amygdala (halo DCN implant)** | Contra: 20<br>Ipsi: 20 | 3 | 9.1<br>14.1 | 1.0<br>2.2 | One-sided Wilcoxon rank sum<br>p=0.140; w=464 |

| | | | | | |
|---|---|---|---|---|---|
| **Amygdala (wildtype)** | 60 total (30/30 contra/ipsi) | 4 | 4.0 | 0.5 | Wilcoxon rank sum<br>vs. contra PBN p=0.57; w=1308<br>vs. ipsi PBN p=3.1e-9; w=2207<br>vs. contra DCN p=8.5e-6; w=1208<br>vs. ipsi DCN p=6.4e-6; w=1213.5 |
| **Basal Forebrain (halo PBN implant)** | Contra: 58<br>Ipsi: 54 | 4 | 4.7<br>11.9 | 0.4<br>0.8 | One-sided Wilcoxon rank sum<br>p = 1.26e-12; w = 2077 |
| **Basal Forebrain (halo DCN implant)** | Contra: 28<br>Ipsi: 28 | 3 | 8.3<br>6.6 | 0.9<br>0.8 | Wilcoxon rank sum<br>p=0.1313; w=706 |
| **Basal Forebrain (wildtype)** | 48 total (24/24 contra/ipsi) | 4 | 7.0 | 0.5 | Wilcoxon rank sum<br>vs. contra PBN p=0.0015; w=2605<br>vs. ipsi PBN p=1.95e-6; w=3489<br>vs. contra DCN p=0.269; w=1180.5<br>vs. ipsi DCN p=0.49; w=1015 |
| **Cingulate Cortex (halo PBN implant)** | Contra: 22<br>Ipsi: 24 | 4 | 29.9<br>47.3 | 6.1<br>4.2 | One-sided Wilcoxon rank sum<br>p = 0.0069; w = 404.5 |
| **Cingulate Cortex (halo DCN implant)** | Contra: 22<br>Ipsi: 22 | 3 | 18.0<br>15.5 | 2.4<br>2.3 | Wilcoxon rank sum<br>P=0.49; w=465 |
| **Cingulate Cortex (wildtype)** | 48 total (24/24 contra/ipsi) | 4 | 32.4 | 1.5 | Wilcoxon rank sum<br>vs. contra PBN p=0.99; w=2605<br>vs. ipsi PBN p=9.75e-7; w=3489<br>vs. contra DCN p=2.7e-5; w=449<br>vs. ipsi DCN p=1.28e-6; w=398 |
| **Piriform Cortex (halo PBN implant)** | Contra: 25<br>Ipsi: 25 | 4 | 17.0<br>31.8 | 2.3<br>3.0 | One-sided Wilcoxon rank sum<br>p=3.79e-4; w = 463.5 |
| **Piriform Cortex (halo DCN implant)** | Contra: 24<br>Ipsi: 24 | 3 | 14.0<br>11.5 | 1.2<br>1.1 | Two-sided Wilcoxon rank sum<br>p=0.134; w=515 |
| **Piriform Cortex (wildtype)** | 48 total (24/24 contra/ipsi) | 4 | 12.6 | 0.5 | Wilcoxon rank sum<br>vs. contra PBN p=0.44; w=991<br>vs. ipsi PBN p=7.04e-8; w=1388<br>vs. contra DCN p=0.19; w=986<br>vs. ipsi DCN p=0.27; w=783.5 |

| | | | | | |
|---|---|---|---|---|---|
| **Preoptic Hypothalamus (halo PBN implant)** | Contra: 16<br>Ipsi: 16 | 4 | 5.9<br>15.3 | 0.7<br>2.0 | One-sided Wilcoxon rank sum<br>p=6.62e-5; w = 162.5 |
| **Preoptic Hypothalamus (halo PBN implant)** | Contra: 24<br>Ipsi: 24 | 3 | 19.0<br>19.1 | 2.4<br>3.1 | Two-sided Wilcoxon rank sum<br>p=0.6; w=614 |
| **Preoptic Hypothalamus (wildtype)** | 32 total (16/16 contra/ipsi) | 4 | 11.7 | 0.8 | Wilcoxon rank sum<br>vs. contra PBN p=0.004; w=378.5<br>vs. ipsi PBN p=0.02; w=654<br>vs. contra DCN p=0.02; w=829<br>vs. ipsi DCN p=0.18; w=765.5 |
| **Anterior & Lateral Hypothalamus (halo PBN implant)** | Contra: 16<br>Ipsi: 16 | 4 | 13.3<br>35.1 | 2.0<br>4.3 | One-sided Wilcoxon rank sum<br>p=1.58e-4; w=168 |
| **Anterior & Lateral Hypothalamus (halo DCN implant)** | Contra: 24<br>Ipsi: 24 | 3 | 17.3<br>15.7 | 1.5<br>1.4 | Two-sided Wilcoxon rank sum<br>p=0.32; w=636.5 |
| **Anterior & Lateral Hypothalamus (wildtype)** | 44 total (22/22 contra/ipsi) | 4 | 9.1 | 0.6 | Wilcoxon rank sum<br>vs. contra PBN p=0.11; w=584.5<br>vs. ipsi PBN p=1.25e-7; w=804<br>vs. contra DCN p4.15e-5; w=1147<br>vs. ipsi DCN p=6.39e-6; w=1179 |
| **Extended Data Figure 11:** | | | | | |
| **Motor Thalamus (halo PBN implant)** | Contra: 18<br>Ipsi: 18 | 3 | 0.6<br>0.5 | 0.4<br>0.3 | One-sided Wilcoxon rank sum<br>p=0.63; w=339.5 |
| **Motor Thalamus (halo DCN implant)** | Contra: 18<br>Ipsi: 18 | 3 | 3.8<br>3.8 | 1.2<br>0.9 | One-sided Wilcoxon rank sum<br>p=0.69; w=347.5 |
| **Motor Thalamus (wildtype)** | 48 total (24/24 contra/ipsi) | 3 | 0.2 | 0.1 | Wilcoxon rank sum<br>vs. contra PBN p=0.29; w=649<br>vs. ipsi PBN p=0.56; w=627.5<br>vs. contra DCN p=1.56e-6; w=865<br>vs. ipsi DCN p=1.25e-5; w=836.5 |

| | | | | | |
|---|---|---|---|---|---|
| **Red Nucleus (halo PBN implant)** | Contra: 18<br>Ipsi: 18 | 3 | 4.2<br>1.6 | 1.0<br>0.4 | One-sided Wilcoxon rank sum<br>p=0.99; w=261.5 |
| **Red Nucleus (halo DCN implant)** | Contra: 22<br>Ipsi: 22 | 3 | 3.1<br>3.1 | 0.6<br>0.4 | One-sided Wilcoxon rank sum<br>p=0.3089; w=516.5 |
| **Red Nucleus (wildtype)** | 36 total (18/18 contra/ipsi) | 3 | 1.8 | 0.4 | Wilcoxon rank sum<br>vs. contra PBN p=0.015; w=625<br>vs. ipsi PBN p=0.939; w=490.5<br>vs. contra DCN p=0.038; w=776<br>vs. ipsi DCN p=0.0074; w=813.5 |
| **Ventral Tegmental Area (halo PBN implant)** | Contra: 18<br>Ipsi: 18 | 3 | 4.5<br>13.3 | 1.0<br>2.4 | One-sided Wilcoxon rank sum<br>p=0.003; w=221 |
| **Ventral Tegmental Area (halo DCN implant)** | Contra: 18<br>Ipsi: 18 | 3 | 22.1<br>21.9 | 2.7<br>3.0 | Two-sided Wilcoxon rank sum<br>p=0.96; w=331 |
| **Ventral Tegmental Area (wildtype)** | 36 total (18/18 contra/ipsi) | 3 | 7.1 | 0.8 | Wilcoxon rank sum<br>vs. contra PBN p=0.01; w=336<br>vs. ipsi PBN p=0.055; w=599.5<br>vs. contra DCN p=5.47e-6; w=742.5<br>vs. ipsi DCN p=2.56e-6; w=751 |

**Table 2: Heart Rate Statistics**

| Figure | Animals | Comparison | Mean Norm. Evoked HR | S.E.M. | Statistics |
|---|---|---|---|---|---|
| **Figure 3G+H:** | | | | | |
| **DCN** | 7 | Evoked Heart rate vs. 1 | 1.12 | 0.08 | Wilcoxon signed rank p = 0.0078; w = 28; one-tailed |
| **PBN** | 7 | Evoked Heart rate vs. 1 | 1.02 | 0.01 | p = 0.078; w = 23; one-tailed |

**Table 3: Place Preference Statistics**

| Figure | Animals | Comparison | Mean | S.E.M. | Statistics |
|---|---|---|---|---|---|
| **Figure 3:** | | | | | |
| **PBN** | 7 | D1 vs<br>D2 | 0.53<br>0.36 | 0.02<br>0.03 | Wilcoxon signed rank<br>p = 0.0078; w = 28; one-tailed |
| | | D2 vs<br>D3 | 0.40 | 0.03 | p = 0.58; w = 10; two-tailed |
| | | D3 vs<br>D4 | 0.50 | 0.03 | p = 0.0078; w = 0; one-tailed |
| | | D2 vs<br>D4 | | | p = 0.0078; w = 0; one-tailed |
| **DCN** | 9 | D1 vs<br>D2 | 0.47<br>0.68 | 0.02<br>0.06 | p = 0.002; w = 0; one-tailed |
| | | D2 vs<br>D3 | 0.54 | 0.04 | p = 0.1641; w = 35; two-tailed |
| | | D3 vs<br>D4 | 0.38 | 0.07 | p = 0.1016; w = 11; one-tailed |
| | | D2 vs<br>D4 | | | p = 0.0098; w = 42; one-tailed |
| **Wildtype** | 9 | D1 vs<br>D2 | 0.51<br>0.48 | 0.02<br>0.04 | p = 0.65; w = 27; two-tailed |

| | | | | | |
|---|---|---|---|---|---|
| | | D2 vs D3 | 0.46 | 0.04 | p = 0.82; w = 25; two-tailed |
| | | D3 vs D4 | 0.52 | 0.04 | p = 0.36; w = 14; two-tailed |
| **Figure 3** | | | | | |
| **PBN:** | | | | | Wilcoxon signed rank |
| D1 | 7 | Vs. chance (0.5) | 0.53 | 0.02 | p = 0.14; w = 21; two-tailed |
| D2 | 7 | | 0.36 | 0.03 | p = 0.0078; w = 0; one-tailed |
| D3 | 7 | | 0.40 | 0.03 | p = 0.0156; w = 1; one-tailed |
| D4 | 7 | | 0.50 | 0.03 | p = 1; w = 14; two-tailed |
| | | | | | Wilcoxon rank sum |
| D1 | 7 | Vs. wildtype | | | p = 0.47; w = 67; two-tailed |
| D2 | 7 | | | | p = 0.02; w = 40; one-tailed |
| D3 | 7 | | | | p = 0.14; w = 49; one-tailed |
| D4 | 7 | | | | p = 0.61; w = 54; two-tailed |
| **DCN:** | | | | | Wilcoxon signed rank |
| D1 | 9 | Vs. chance (0.5) | 0.47 | 0.02 | p = 0.3008; w = 13; two-tailed |
| D2 | 9 | | 0.68 | 0.06 | p = 0.0098; w = 42; one-tailed |
| D3 | 9 | | 0.54 | 0.04 | p = 0.1797; w = 31; one-tailed |
| D4 | 9 | | 0.38 | 0.07 | p = 0.4961; w = 16; two-tailed |
| | | | | | Wilcoxon rank sum |
| D1 | 9 | Vs. wildtype | | | p = 0.3865; w = 75; two-tailed |
| D2 | 9 | | | | p = 0.0313; w = 107; one-tailed |
| D3 | 9 | | | | p = 0.068; w = 103; one-tailed |
| D4 | 9 | | | | p = 0.2973; w = 73; two-tailed |
| **Wt:** | | | | | Wilcoxon signed rank |
| D1 | 9 | Vs. chance (0.5) | 0.51 | 0.02 | p = 0.82; w = 20; two-tailed |
| D2 | 9 | | 0.48 | 0.04 | p = 0.91; w = 21; two-tailed |
| D3 | 9 | | 0.46 | 0.04 | p = 0.43; w = 15; two-tailed |
| D4 | 9 | | 0.52 | 0.04 | p = 0.65; w = 27; two-tailed |
| **Extended Data Figure 8:** | | | | | |
| **PBN (unilateral)** | 5 | D1 vs D2 | 0.50<br>0.37 | 0.04<br>0.05 | Wilcoxon signed rank<br>p = 0.0313; w = 15; one-tailed |

| | | | | | |
|---|---|---|---|---|---|
| | | D2 vs D3 | 0.43 | 0.06 | p = 0.625; w = 5; two-tailed |
| | | D3 vs D4 | 0.37 | 0.29 | p = 0.906; w = 12; one-tailed |
| | | D4 vs D2 | | | p = 0.7813; w = 10; one-tailed |
| **DCN (unilateral)** | 3 | D1 vs D2 | 0.49<br>0.73 | 0.05<br>0.16 | p = 0.125; w = 0; one-tailed |
| | | D2 vs D3 | 0.42 | 0.05 | p = 0.250; w = 6; two-tailed |
| | | D3 vs D4 | 0.51 | 0.09 | p = 1; w = 0; one-tailed |
| | | D4 vs D2 | | | p = 0.125; w = 0; one-tailed |
| **Extended Data Figure 8:** | | | | | |
| **PBN (unilateral):** | | | | | Wilcoxon signed rank |
| D1 | 5 | Vs. chance (0.5) | 0.50 | 0.04 | p = 1; w = 8; two-tailed |
| D2 | 5 | | 0.37 | 0.05 | p = 0.063; w = 1; one-tailed |
| D3 | 5 | | 0.43 | 0.06 | p = 0.156; w = 3; one-tailed |
| D4 | 5 | | 0.37 | 0.29 | p = 0.063; w = 0; two-tailed |
| | | | | | Wilcoxon rank sum |
| D1 | 5 | Vs. wildtype | | | p = 1; w = 38; two-tailed |
| D2 | 5 | | | | p = 0.095; w = 27; one-tailed |
| D3 | 5 | | | | p = 0.303; w = 33; one-tailed |
| D4 | 5 | | | | p = 0.042; w = 22; two-tailed |
| **DCN (unilateral):** | | | | | Wilcoxon signed rank |
| D1 | 3 | Vs. chance (0.5) | 0.49 | 0.05 | p = 1; w = 3; two-tailed |
| D2 | 3 | | 0.73 | 0.11 | p = 0.125; w = 6; one-tailed |
| D3 | 3 | | 0.42 | 0.05 | p = 1; w = 0; one-tailed |
| D4 | 3 | | 0.51 | 0.09 | p = 1; w = 3; two-tailed |
| | | | | | Wilcoxon rank sum |
| D1 | 3 | Vs. wildtype | | | p = 0.482; w = 15; two-tailed |
| D2 | 3 | | | | p = 0.050; w = 29; one-tailed |
| D3 | 3 | | | | p = 0.759; w = 16; one-tailed |
| D4 | 3 | | | | p = 1; w = 19; two-tailed |

**Table 4: Open Field Statistics**

| Figure | Animals | Comparison | Mean | S.E.M. | Statistics |
|---|---|---|---|---|---|
| **Open Field – Extended Data Figure 7** | | | | | |
| Time in Center | | | | | |
| **Wildtype** | 11 | Laser Off vs. On | 0.11<br>0.08 | 0.02<br>0.02 | Wilcoxon Signed rank<br>p = 0.067; w = 54; two-tailed |
| **DCN** | 7 | Laser Off vs. On | 0.10<br>0.11 | 0.03<br>0.04 | p = 0.948; w = 15; two-tailed |
| **PBN** | 6 | Laser Off vs. On | 0.14<br>0.12 | 0.04<br>0.05 | p = 0.094; w = 19; two-tailed |
| Average Velocity | | | | | |
| **Wildtype** | 11 | Laser Off vs. On | 6.13<br>6.18 | 0.53<br>0.58 | p = 0.97; w = 32; two-tailed |
| **DCN** | 7 | Laser Off vs. On | 5.10<br>4.43 | 0.60<br>0.59 | p = 0.02; w = 28; two-tailed |
| **PBN** | 6 | Laser Off vs. On | 5.07<br>4.93 | 0.43<br>0.49 | p=0.44; w = 15; two-tailed |

| Clust# | # cells | Relevant markers | % in cluster w/ marker | % of PBN cells | Putative Location | Evidence/Validation |
|---|---|---|---|---|---|---|
| 0 | 576 | Unassigned | | | | |
| 1 | 472 | Foxp2 | 28.8% | 29.4% | Parabrachial | Figure 5 immunohistochemistry<br>Allen Brain Atlas in situ |
| 2 | 342 | Lmx1a | 51% | 21.3% | Parabrachial | Figure 5 in situ |
| 3 | 295 | Lamp5 | 42% | - | Vestibular & Tegmentum (PDTG) | Allen Brain Atlas in situ |
| 4 | 248 | Sox2;<br>Crhbp | 30%<br>24% | - | Vestibular | Allen Brain Atlas in situ |
| 5 | 212 | Grp<br>Tacr1 | 67%<br>29% | 13.2% | Parabrachial | Allen Brain Atlas in situ<br>Allen Brain Atlas in situ |
| 6 | 207 | Dbh;<br>TH | 88%<br>85% | - | Locus Coeruleus | Allen Brain Atlas in situ<br>Allen Brain Atlas in situ |
| 7 | 190 | Necab1<br>Pax5<br>Tacr1 | 78%<br>33.2%<br>23.2% | 11.8% | Parabrachial | Figure 5 in situ<br>Allen Brain Atlas in situ<br>Allen Brain Atlas in situ |
| 8 | 135 | Penk;<br>Tacr1<br>Pax5<br>Brs3 | 73%<br>48%<br>43%<br>24% | 8.4% | Parabrachial | Allen Brain Atlas in situ<br>Allen Brain Atlas in situ<br>Figure 5 in situ<br>Allen Brain Atlas in situ |
| 9 | 132 | Gabra6 | 70.5% | - | Cerebellar granule cell | Allen Brain Atlas in situ |
| 10 | 127 | Pax5 | 70% | 7.9% | Parabrachial | Figure 5 in situ |
| 11 | 127 | Slc6a5<br>Pde3a | 50%<br>39.4% | 7.9% | Parabrachial | Herbert et al., 2000[1]<br>Allen Brain Atlas in situ |
| 12 | 109 | Crh | 57% | - | Barrington's | Allen Brain Atlas in situ |
| 13 | 101 | Chat<br>Slc5a7 | 86%<br>89% | - | PDTG Cholinergic neurons | Allen Brain Atlas in situ<br>Allen Brain Atlas in situ |
| 14 | 99 | Lhx4 | 21.2% | - | Tegmentum (PDTG) | Allen Brain Atlas in situ |
| 15 | 98 | Lmx1a | 58% | - | PBN | Figure 5 in situ |
| 16 | 80 | Kit<br>PCP2 | 70%;<br>33% | - | Cerebellar gabaergic neurons | Allen Brain Atlas in situ<br>Allen Brain Atlas in situ |
| 17 | 72 | Piezo2 | 99% | - | Mesencephalic Trigeminal | Florez-Pax et al., 2016[2] |
| 18 | 66 | Penk<br>Lmx1a<br>Drd1 | 89%<br>54.5%<br>35% | - | PBN | Allen Brain Atlas in situ<br>Figure 5 in situ<br>Allen Brain Atlas in situ |
| 19 | 55 | Crhbp | 71% | - | Vestibular | Allen Brain Atlas in situ |

| | | | | | | |
|---|---|---|---|---|---|---|
| 20 | 48 | Col27a1 | 63% | - | Dorsal Cochlear | Allen Brain Atlas in situ |
| 21 | 45 | Eomes | 40% | - | Cerebellar unipolar brush cell | Allen Brain Atlas in situ |
| 22 | 25 | PCP2 | 28% | - | Cerebellar Purkinje cell | Allen Brain Atlas in situ |
| 23 | 15 | Contamination (Gfap) | 73% | - | Glia | |

**Table 5: RNAseq Cluster Localization.**

The expression of the noted genes and references noted in the right-most column were used to categorize the different clusters in **Figure 5**. Different shading is used to indicate putative PBN clusters (*blue*), cerebellar clusters and clusters with fewer than 100 neurons (*grey*), and clusters likely outside of the PBN (*white*).

| Gene | Cre-line used | Location in PBN | Observed projections in forebrain | Observed behaviors | Clusters |
|---|---|---|---|---|---|
| Foxp2 | Foxp2-IRES-Cre (Jax 030541)[3] | Across PBN & concentration in anterior/lateral[3,4] | Septum, basal ganglia, amygdala, basal forebrain, hypothalamus, thalamus[3] | Unknown | 1, 5, 8, 11 |
| Lmx1a | Tg(Lmx1a-cre)1Kjmi[4,5] | Across posterior PBN & concentration in anterior/lateral[4] | Unknown | Unknown | 2 |
| Grp | None (FISH used[4]) | Dorsal “head” and “waist” PBN[4] | Unknown | Unknown | 5 |
| Satb2 | Satb2-IRES-Cre[6] (Jax 030546) | Across PBN, waist PBN[4] | Cortex, amygdala, thalamus[6,7] | Taste preference[6] | 2 |
| Tacr1 | Tacr1-T2A-Cre-Neo[8] (Allen Institute) | Across PBN, concentrated in dorsal[8] | Hypothalamus, thalamus[8] | Necessary and sufficient to drive pain-related behaviors[8] | 5, 7, 8 |
| Penk | Penk-IRES2-Cre[9] (Jax 025112) | Across PBN & concentration in anterior-lateral PBN[9] | Hypothalamus[9] | Place aversion, thermoregulation[9] | 1, 2, 5, 8 |
| Brs3 | Brs3-resCRE:GFP[10] (Jax 030540) | Across PBN, only lateral PBN described[10] | Amygdala, Thalamus, Hypothalamus , VTA[10] | Unknown | 8 |
| Slc6a5 (Glyt2) | None (IHC)[1] | Medial PBN, Kolliker-Fuse[1] | Unknown | Unknown | 11 |
| Slc17a6 (vGlut2) | Slc17a6-IRES-Cre (Jax 016963)[3] | Almost all PBN neurons[3,4] | Cortex, Septum, Amygdala, Basal Forebrain, Hypothalamus, Thalamus[3,7] | Place aversion, thermoregulation[9] | 1, 2, 5, 7, 8, 10 |
| Calca (CGRP) | Calca-Cre (Jax 033168)[11,12] | Anterior + Lateral PBN[11] | Cortex, Septum, Basal ganglia, Amygdala, Basal forebrain, Thalamus[11] | General alarm[12] | None |
| Pdyn | Pdyn-IRES-Cre (Jax 027958)[3,4,9] | Lateral PBN + Pre-Coeruleus[3,4,9] | Hypothalamus, Thalamus[3,9] | Place aversion, thermoregulation[9] | None |

**Table 6: Cluster markers and their association with different behaviors.** A number of the genes identified in the RNAseq experiments of **Figure 5** are known markers of the PBN. Previous studies have used the indicated cre lines to study the anatomical projections and behavioral roles of different subtypes of PBN neurons.  Listed are relevant genes the cre lines used, their locations within the PBN and the observed projections in the forebrain as well as their roles in different behaviors.  In some cases a single gene identifies a single cluster, whereas in other cases multiple genes correspond to multiple clusters.

| | PC→PBN pathway | PC→LC pathway |
|---|---|---|
| **Fig. 2 g, h** | High density of PC-PBN synapses (40X as many PC-PBN synapses as PC-LC synapses) | Very low density of PC synapses |
| **Fig. 2 d-f, i** | Shows extensive PC synapses throughout the PBN. | - |
| **Fig. 2 k.** | Optogenetic suppression of PC firing led to short latency firing in 50% of PBN neurons *in vivo*. This shows that PCs in the posterior vermis inhibit a large fraction of PBN neurons. | - |
| **Fig. 2 l, m.** | Optogenetic stimulation of PC fibers in brain slice evoked large IPSCs in 75% of PBN neurons with a latency of 2.3 ms. This establishes that PCs directly inhibit a large fraction of neurons in the posterior PBN neurons. | - |
| **Fig. 3 c-f.** | PBN neurons are selectively activated in behavior studies targeting the PBN pathway. | LC cells are not activated in behavioral experiments targeting the PBN pathway. |
| **Extended Data Fig. 9.** | Shows labeled neurons for the 3 mice used for anatomical studies. Fluorescence is localized to the PBN, indicating that axonal labelling in the rest of the panel and in Figure 4 is due to the PBN | Shows labeled neurons for the 3 mice used for anatomical studies. There are no fluorescently labelled cells in the LC for any of the mice, indicating that the fluorescence labelling in Figure 4 is not a result of LC labelling. |
| **Fig. 5 a-c** | Identification of multiple types of PBN neurons targeted by PCs. Readily interpretable because the PBN does not project back to the cerebellum. [3,11] | AAV1-Syn-Cre labelling of LC neurons is not interpretable because LC neurons project extensively to the cerebellum and could be retrogradely labelled[13-15]. This is supported by the observation that many of the neurons labelled by AAV1-Syn-Cre in the LC are not located in the vicinity of PC fibers. |
| **Fig. 5e.** | Shows that trans-synaptically labelled neurons within the PBN are in the vicinity of PC fibers. | |
| **Fig. 5g** | Shows that there are neurons within the PBN that are directly inhibited by PCs that in turn project to the amygdala. | |
| **Fig. 5 h-j.** | Establishes the identity of some types of PBN neurons targeted by PCs. These neurons have been studied previously. There projection pattern overlaps with the projection pattern in our trans-synaptic studies, and place preference studies indicate that they are aversive, as in our behavioral studies. | - |
| **Schwarz et al., *Nature* 2015. Figure S6[15]** | - | This figure shows that PC fibers are only present at the very posterior/dorsal edges of the LC, and that PC synapses within the LC are rare. |

**Table 7: Summary of the evidence supporting the PC-PBN pathway rather than the PC-LC pathway in our experimental results.** It was important to consider whether the PC to LC pathway could contribute to some of our experimental findings in light of the close proximity of the LC to the PBN, and a previous report of a PC to LC pathway [15]. Our experiments were performed with that possibility in mind, and as summarized in this table, our results establish that the PC-PBN pathway is the dominant contributor.

**Extended Data References**


1 Herbert, H., Guthmann, A., Zafra, F. & Ottersen, O. P. Glycine, glycine receptor subunit and glycine transporters in the rat parabrachial and Kolliker-Fuse nuclei. *Anat Embryol (Berl)* **201**, 259-272, doi:10.1007/s004290050316 (2000).

2 Florez-Paz, D., Bali, K. K., Kuner, R. & Gomis, A. A critical role for Piezo2 channels in the mechanotransduction of mouse proprioceptive neurons. *Sci Rep* **6**, 25923, doi:10.1038/srep25923 (2016).

3 Huang, D., Grady, F. S., Peltekian, L. & Geerling, J. C. Efferent projections of Vglut2, Foxp2, and Pdyn parabrachial neurons in mice. *J Comp Neurol* **529**, 657-693, doi:10.1002/cne.24975 (2021).

4 Karthik, S. *et al.* Molecular ontology of the parabrachial nucleus. *J Comp Neurol* **530**, 1658-1699, doi:10.1002/cne.25307 (2022).

5 Chizhikov, V. V. *et al.* The roof plate regulates cerebellar cell-type specification and proliferation. *Development* **133**, 2793-2804, doi:10.1242/dev.02441 (2006).

6 Jarvie, B. C., Chen, J. Y., King, H. O. & Palmiter, R. D. Satb2 neurons in the parabrachial nucleus mediate taste perception. *Nat Commun* **12**, 224, doi:10.1038/s41467-020-20100-8 (2021).

7 Grady, F., Peltekian, L., Iverson, G. & Geerling, J. C. Direct Parabrachial-Cortical Connectivity. *Cereb Cortex* **30**, 4811-4833, doi:10.1093/cercor/bhaa072 (2020).

8 Barik, A. *et al.* A spinoparabrachial circuit defined by Tacr1 expression drives pain. *Elife* **10**, doi:10.7554/eLife.61135 (2021).

9 Norris, A. J., Shaker, J. R., Cone, A. L., Ndiokho, I. B. & Bruchas, M. R. Parabrachial opioidergic projections to preoptic hypothalamus mediate behavioral and physiological thermal defenses. *Elife* **10**, doi:10.7554/eLife.60779 (2021).

10 Mogul, A. S. *et al.* Cre Recombinase Driver Mice Reveal Lineage-Dependent and -Independent Expression of Brs3 in the Mouse Brain. *eNeuro* **8**, doi:10.1523/ENEURO.0252-21.2021 (2021).

11 Huang, D., Grady, F. S., Peltekian, L., Laing, J. J. & Geerling, J. C. Efferent projections of CGRP/Calca-expressing parabrachial neurons in mice. *J Comp Neurol* **529**, 2911-2957, doi:10.1002/cne.25136 (2021).

12 Palmiter, R. D. The Parabrachial Nucleus: CGRP Neurons Function as a General Alarm. *Trends Neurosci* **41**, 280-293, doi:10.1016/j.tins.2018.03.007 (2018).

13 Loughlin, S. E., Foote, S. L. & Bloom, F. E. Efferent projections of nucleus locus coeruleus: topographic organization of cells of origin demonstrated by three-dimensional reconstruction. *Neuroscience* **18**, 291-306, doi:10.1016/0306-4522(86)90155-7 (1986).

14 Steindler, D. A. Locus coeruleus neurons have axons that branch to the forebrain and cerebellum. *Brain Res* **223**, 367-373, doi:10.1016/0006-8993(81)91149-5 (1981).

15 Schwarz, L. A. *et al.* Viral-genetic tracing of the input-output organization of a central noradrenaline circuit. *Nature* **524**, 88-92, doi:10.1038/nature14600 (2015).
